# A Caveat Regarding the Unfolding Argument: Implications of Plasticity

**DOI:** 10.1101/2025.11.04.686457

**Authors:** Vikas N. O’Reilly-Shah, Alessandro Maria Selvitella, Aaron Schurger

## Abstract

The unfolding argument in the neuroscience of consciousness posits that causal structure cannot account for consciousness because any recurrent neural network (RNN) can be “unfolded” into a functionally equivalent feedforward neural network (FNN) with identical input-output behavior. Subsequent debate has focused on dynamical properties and philosophy of science critiques. We examine a boundary condition on the unfolding argument for RNN systems with rapid plasticity in their connection weights. We demonstrate through rigorous mathematical proofs that rapid plasticity negates the functional equivalence between RNN and FNN. Our proofs address history-dependent plasticity, dynamical systems analysis, information-theoretic considerations, perturbational stability, complexity growth, and resource limitations. We demonstrate that neuronal systems that possess properties such as plasticity, history-dependence, and complex temporal information encoding have features that cannot be captured by a static FNN. We show that plasticity is a concrete instance of lenient dependency between behavioral and internal observables, restoring empirical testability to theories that incorporate plasticity on perception-relevant timescales. Our results do not establish that recurrence, plasticity, or process are necessary for consciousness; they establish that the unfolding argument does not preclude empirical investigation of whether these properties matter.

## Introduction

Despite significant advances in both neuroscience and computation, a scientific explanation of consciousness remains elusive (Seth & Bayne 2022). Modern physicalist theories of consciousness, in particular integrated information theory (IIT), have focused on the causal structure of neural network systems (Albantakis et al. 2023). Recurrent causal structures are necessary for a system to have cause-effect power upon itself, an axiomatic principle in IIT for the development of consciousness from such a system. Among the many challenges facing physicalist theories of consciousness, the so-called *unfolding argument* (UA) poses a significant obstacle to any theory that asserts recurrence as a fundamental requirement.

### The Unfolding Argument

Doerig and colleagues (2021; 2019) developed the unfolding argument as a challenge to causal structure theories of consciousness - theories that identify consciousness with specific patterns of causal interaction rather than input-output behavior. Their argument targets theories like Integrated Information Theory (IIT) and Recurrent Processing Theory (RPT), which propose that consciousness depends on how neural elements interact with each other, not merely on what function they compute.

The unfolding argument rests on four key premises, paraphrased from the original paper (Doerig et al. 2019):

- **P1**: Scientific investigation must rely on physical measurements, and in the investigation of consciousness these are all ultimately forms of subjective report
- **P2**: Any recurrent neural network (RNN) can be "unfolded" into a functionally equivalent feedforward neural network (FNN)
- **P3**: Systems with identical input-output functions cannot be distinguished by experiments that rely on physical measurements, except for measurements of the internal workings of the system - i.e. brain activity
- **P4**: We cannot use measures of brain activity as *a priori* indicators of consciousness since understanding the brain basis of consciousness is what we seek to explain in the first place.

From these premises, Doerig et al. (2019) conclude that causal structure theories face a fatal dilemma: they are either false (if they accept that unfolded feedforward networks can be conscious) or unscientific (if they maintain that functionally identical systems with different causal structures have different consciousness despite being empirically indistinguishable).

The unfolding argument has faced significant criticism since it was introduced. Critiques have challenged its second premise, arguing that the claimed functional equivalence is not robust to dynamic or structural perturbations (Usher 2021). Others have questioned its philosophical basis, asserting that its restrictive, behaviorist-like methodology is inappropriate for consciousness science (Usher et al. 2023). Kleiner and Hoel (2021) generalized this challenge into a broader "substitution argument," demonstrating that the independence of a theory’s predictive data (internal observables) from its inferential data (reports) leads to the *a priori* falsification of any minimally informative theory. This generalization has prompted further debate, with critics arguing that certain classes of functionalist theories are immune to such substitutions (Kleiner & Hoel 2021; Ganesh 2020). Notably, in an article responding to various critiques, Herzog, Schurger, and Doerig (2022) claim an extension of the unfolding argument to all functions that Turing Complete (TC) systems can calculate, and therefore "any function, and even every process can be implemented with many different causal structures." This is a significant generalization. Understood in the broader context, the unfolding argument and its Turing Completeness generalization can be seen as technical instantiations of the classical multiple realizability (MR) argument from philosophy of mind (Shapiro 2000; Kim 1992). MR, as originally formulated by Putnam (1967), holds that the same mental state type can be realized by systems with different physical or causal structures. The UA makes this claim precise for neural networks; the TC generalization extends it to arbitrary computational substrates.

However, critical gaps in the literature exist. First, there has been a failure to rigorously define the systems under consideration (FNN and RNN) from the outset. Neither the Doerig paper nor any subsequent paper has made any attempt to mathematically formalize the argument, relying instead on references to the computer science and neural network literature. Second, there has been a failure to consider how synaptic plasticity might alter the functional properties of a recurrent neural network.

Therefore, in this paper, we explore how the non-stationarity introduced by plasticity in a recurrent neural network results in a system that cannot be durably replicated by any static feedforward architecture. Plasticity helps to address the Kleiner-Hoel challenge. Our technically grounded argument provides a critical caveat to the unfolding argument and establishes a new set of theoretical bounds on the testability of theories of consciousness.

### The Plasticity Caveat

It is well understood that biological neural networks do not merely process static inputs; the dynamical trajectory of neuronal activity is key to understanding how they function (Rabinovich et al. 2006). Biological neural networks are not static computational devices but continuously evolving systems where connectivity patterns change based on activity history. This plasticity operates across multiple timescales, including millisecond-scale synaptic facilitation that is relevant to the timescale on which perception unfolds (Chambers & Rumpel 2017; Magee & Grienberger 2020; Fauth & Tetzlaff 2016; Zenke & Gerstner 2017).

To formally explore the implications of plasticity for the unfolding argument and for theories of consciousness, we begin by defining the systems under consideration (S1 Supplementary Appendix, Introduction). We recognize that the biophysics underlying actual neurobiological systems is continuous in nature; the biological systems under consideration are continuously evolving. However, for the purposes of the mathematical discussion, we treat these as discretely evolving systems (systems evolving in discrete time steps). The unfolding argument itself relies on a treatment of neural network systems as discrete (an RNN is unfolded into the equivalent FNN based on the timesteps of the input to the RNN). Beyond that, treatment of these systems as discretely evolving captures the essence of the key considerations associated with the introduction of plasticity, while avoiding complexities necessitated in the treatment of continuously evolving systems.

We proceed to reiterate the unfolding argument using the rigorously defined mathematical systems under consideration (S1 Supplementary Appendix, Introduction). We then mathematically introduce the implications for the unfolding argument when the input-output mapping in the RNN is allowed to change over time (plasticity in the RNN). Formally, let us consider the RNN *R*_D_, which denotes an RNN with plasticity defined as such.

When such plasticity is present, the unfolding argument becomes substantively bounded. Suppose we have the FNN *F*_D_ such that *F*_D_ reflects the unfolded functional equivalent of the RNN *R*_D_ at the time of unfolding (call it *t_0_*). We demonstrate that for all *t* > *t_0_*, the evolution of the input-output map in *R*_D_ results in divergence between *R*_D_ and the originally unfolded FNN *F*_D_. A key distinction here is that the trajectory divergence we are identifying is a divergence in the *function space* - the mapping itself between inputs and outputs is evolving. This is as compared to the *state space* trajectory; a fixed RNN may describe a state space trajectory *within* its fixed weights in a way that a fixed FNN cannot (as pointed out by Usher (2021)). Concretely, a fixed-weight RNN processing a sequence traverses its hidden state attractor but always implements the same function φ, say, recognizing faces. A plastic RNN might start implementing φ_0_ (recognizing faces) but end, after plasticity implementing φ_t_ (recognizing faces and associating learned names). The function itself has changed - movement in function space, not just state space.

#### Claims

To ensure that the claims made in this manuscript are clear, we enumerate them below. These claims are the subject of the analysis presented in and taken up by the remainder of the manuscript.

1. **We do claim that the UA’s premise does not apply to plastic systems, a caveat to UA premise P2.** The UA compares systems with fixed input-output functions *at a single time step*. Plastic systems do not have fixed functions; they traverse function space over time. The question "can a system with different internal organizational structure compute the same function?" presupposes there is a single function to compare. For plastic systems, this presupposition is false, because the internal organization is undergoing temporal evolution. The argument remains relevant for theories that can be reduced to static organizational structures. The Plasticity and the Unfolding Argument section outlines these arguments; S1 Supplementary Appendix Sections 3-7 details them.
2. **We do claim that perturbational differences outlined directly challenge UA premise P3.** This difference is distinct from Usher’s original perturbation argument (Usher 2021); both challenge the unfolding argument, but Usher’s argument operates in the hidden state space trajectory of a fixed RNN, whereas our argument operates in the function space trajectory of a plastic RNN, a more powerful result. Information-theoretic differences also provide a challenge to P3.
3. **We do <u>not</u> claim hypercomputation, or that substrate matters for consciousness.** We agree that the Turing Complete generalization implies that any plastic RNN could be implemented in an alternative substrate. A plastic RNN implemented in silicon with the same dynamical internal organization as one in biological tissue would be functionally indistinguishable on our analysis. However, these two systems might be distinguishable on perturbation. Additionally, in order for the alternative substrate to causally track the behavior of the original, it would need access to the input data stream of the original. Finally, a Turing Machine cannot *a priori* know the trajectory of the plastic RNN without precomputing it. These points are causally relevant to multiple realizability; neither operates outside the computability parameters of the standard Church-Turing thesis.
4. **We do <u>not</u> claim, or advocate for, any specific theory of consciousness.** We present boundary conditions on the UA that may or may not apply to specific theories. Implications for particular formulations are discussed, but this manuscript should not be viewed as advocating for a specific theory.

#### Direct Response to the Unfolding Argument

Our analysis does not refute the original unfolding argument within its stated constraints - namely, functional input-output equivalence over finite time horizons with fixed parameters. Rather, we formally delineate the boundary conditions within which this equivalence holds. This has significant implications for computational theories of consciousness. Specifically, we show that for theories that explicitly incorporate plastic recurrent dynamics, the challenge presented by the unfolding argument is avoided. The argument remains relevant for theories that can be reduced to static causal structures (arguably: (Usher 2021; Usher et al. 2023; Negro 2020; Ganesh 2020)), but loses force when applied to genuinely dynamic, adaptive systems that change their own structure through experience on the same short time scales as the neuronal interactions underlying perceptual decision making.

First, for premise P2, which asserts that any recurrent system can be unfolded into a functionally equivalent feedforward one, we demonstrate that this premise is invalidated under conditions of plasticity. We demonstrate that this claimed equivalence breaks down due to learning-induced functional divergence, the exponential growth of a plastic system’s functional repertoire, and the prohibitive resource scaling required for unfolding.

Similarly, premise P3 is also undermined. This premise claims that functionally identical systems are empirically indistinguishable. This statement is not directly challenged, but rather, we show that the antecedent condition to P3 is not met because we demonstrate that plastic recurrent systems and their static feedforward counterparts are not functionally identical. Our arguments do not involve looking ’inside’ the network, but rather demonstrate that the experimenter can ascertain based on the behavior of the system if it is feedforward vs plastic recurrent. This is because their learning trajectories may diverge over relevant timescales (depending on the learning rate), because they possess different information-theoretic capacities for memory and prediction, and because they exhibit distinguishable responses to perturbation. To distinguish systems empirically, one could conduct controlled experiments where multiple systems are tested under the same perturbation protocol. The key is that the different behavioral patterns emerge from the input-output function itself, not from examining internal architecture.

#### Proofs

We present here a description of the formal arguments that establish precise mathematical boundaries on the applicability of the unfolding argument to RNN systems with plasticity. The formal proofs are detailed in the S1 Supplementary Appendix. The summaries of these arguments, presented below, serve as plain language explanations of these rigorous proofs. A note on terminology as it relates to this section: we use the terms ’learning’ and ’plasticity’ in this section interchangeably and only in the purely mathematical sense. Both terms, for the purposes of this section, refer to a nontrivial change to the network parameters (weights) between timesteps. *Learning*, here, is not meant in the more biological, task-directed sense.

Let us continue to examine an RNN *R*_D_ with an evolving input-output map as a function of time. Let us also continue to consider the unfolded equivalent FNN *F*_D_. As stated, we concede that those networks would reflect an equivalent input-output function at the time of unfolding (call it *t_0_*). While an RNN and an FNN might be functionally equivalent at that initial time point, any non-trivial plasticity in the RNN will alter its parameters and, due to the evolving nature of neural network functions, necessarily change its input-output mapping over time. This evolution in the input-output function is mathematically represented as □*^R^_t_ : X^t^* → Y. Since the FNN is static, this functional equivalence breaks for all *t* > *t_0_* (Figure 1).

**Figure 1:**
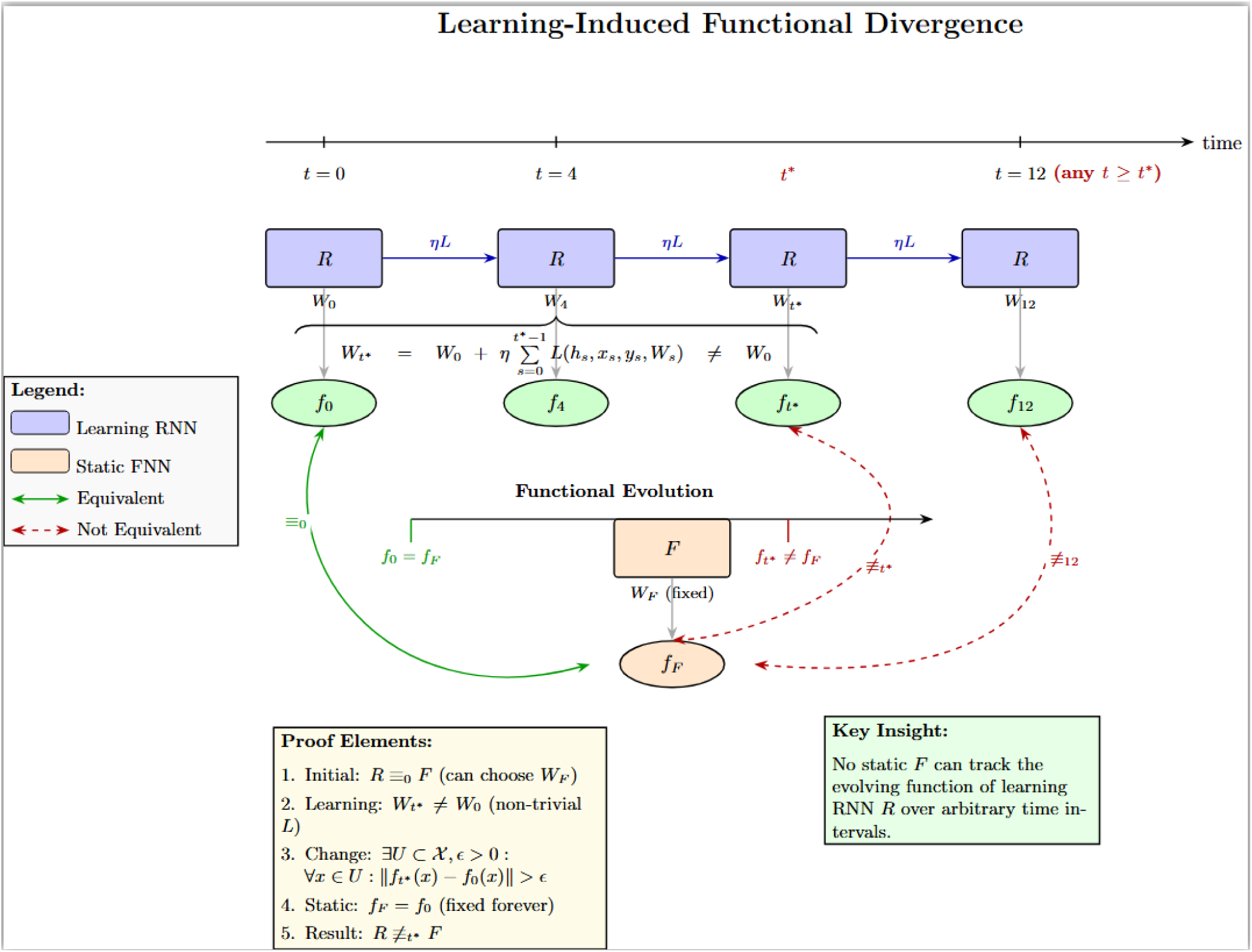
Visual representation of the functional divergence argument. A recurrent neural network *R* with plastic weights evolves through plasticity rule *P*, while a static feedforward network *F* maintains fixed parameters *W_F_*. Initially at *t*=0, both networks can be functionally equivalent, implementing the same input-output mapping *f*_0_=*f_F_*. However, as *R* updates and its parameters evolve, its implemented function changes from *f_0_* to *f*_□□_. Since *F* cannot modify its function *f_F_*, functional equivalence necessarily breaks at some time *t*_□_ > 0 (shown in red). This divergence persists for all subsequent times, demonstrating that no single static network can maintain functional equivalence with a plastic system over arbitrary time intervals.

The key insight is that plasticity, a ubiquitous feature of biological neural systems, inherently creates time-varying functions that are incompatible with any static representation (S1 Supplementary Appendix, Learning Limits Unfolding). In fact, even more fundamentally, a static FNN processes a fixed number of variables as input (corresponding to the matching time *t_0_*), while the input space of the RNN expands to calculate the output with information gathered at each time step through nontrivial weight changes (plasticity). This increases the dimensionality of the input space. Biologically plausible mechanisms like Hebbian learning or spike-timing-dependent plasticity continuously modify synaptic strengths based on neural activity. This directly challenges premise P2 of the unfolding argument; if consciousness depends on systems that learn and adapt, no single static FNN can replicate their behavior over extended periods. We also note that the folding-unfolding relationship is asymmetric: any feedforward network can be trivially folded into a recurrent network, but the converse does not hold in general, establishing strict containment.

Next, we present an information-theoretic analysis, targeting the claim that FNNs can match the information processing capabilities of RNNs (S1 Supplementary Appendix, Learning Implies More Information). The core insight is that the fixed-window architecture of an FNN imposes a hard bound on its memory, while the internal state of the equivalent RNN has a much more robust informational capacity. This is particularly relevant for biological functions like working memory and episodic memory, which require integrating information over extended and variable time periods. We formally prove that, due to the monotonicity of mutual information, a fixed-window FNN is necessarily less informative for prediction tasks than an RNN that can capture this historical information (Figure 2). This challenges premise P2 of the unfolding argument by demonstrating that RNNs and static FNNs can be distinguished by their information processing capacities, especially in tasks requiring long-term memory.

**Figure 2:**
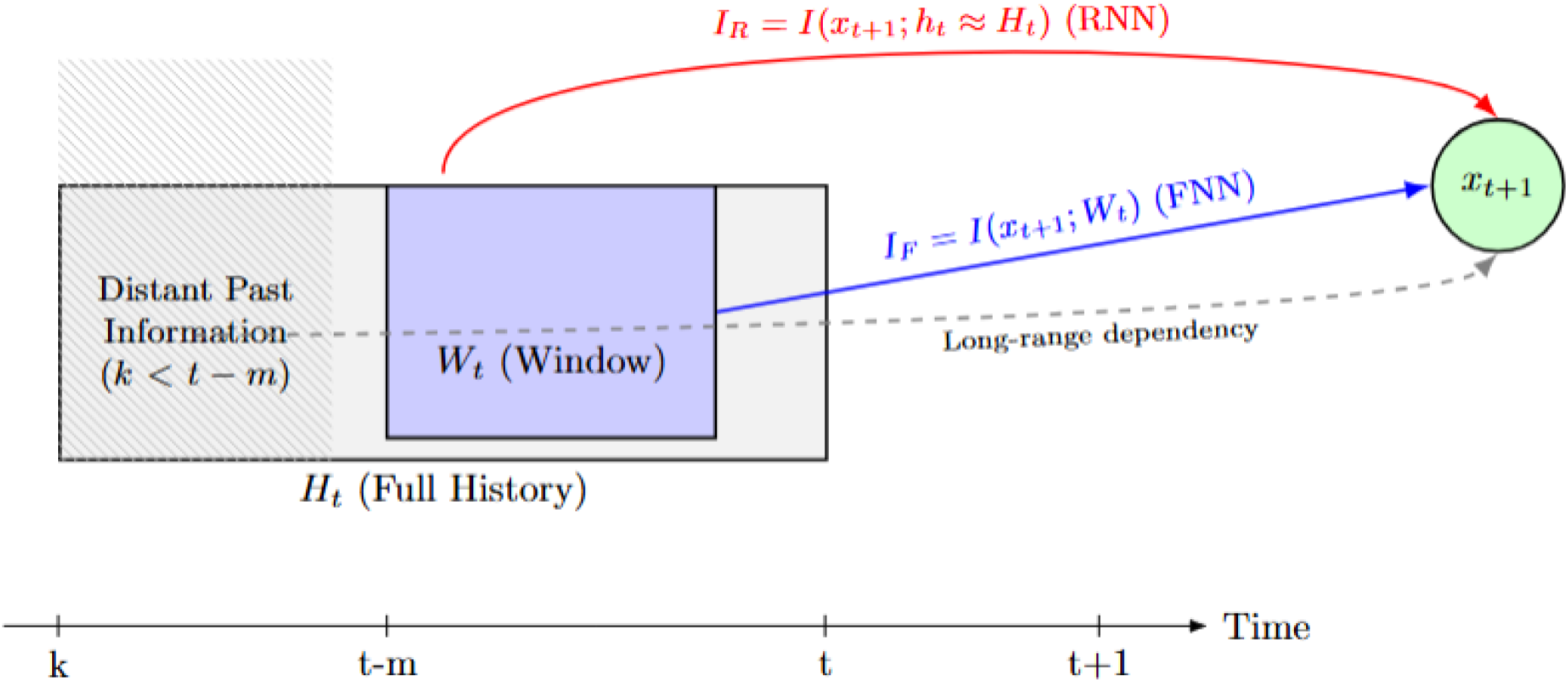
Visual representation of the argument from information-theoretic limitations. The complete history of the process up to time *t* is represented by *H_t_*(large gray box), which includes both recent past and distant past information (hatched area, *k* < *t-m*). A static Feedforward Neural Network (FNN) predicts the next state *x_t+1_* (green circle) using only information from a fixed recent window *W_t_* (blue box), capturing predictive information *I_F_* (blue arrow). A Recurrent Neural Network (RNN), through its internal state *h_t_*, can potentially summarize the entire history *H_t_* and uses this to predict *x_t+1_*, capturing predictive information *I_R_*(red arrow). There is non-zero information from the distant past (hatched area) relevant for predicting *x_t+1_* (represented by the dashed gray arrow indicating a long-range dependency). This information is available to the RNN but not to the FNN, which is limited to the window *W_t_*. Consequently, the FNN’s predictive capacity is strictly limited compared to the RNN’s (*I_F_* < *I_R_*).

We then examine the fundamental non-equivalence from the perspective of dynamical systems theory (S1 Supplementary Appendix, Learning at Discrete Times Needs Countable Unfolding). This examination targets the assumption that an FNN can capture the essential dynamical properties of an RNN. The critical insight here is that RNNs are orbit generators capable of long-term temporal evolution, whereas FNNs compute finite-horizon maps (Figure 3). When the NNs are considered from the perspective of the function space they represent, a static FNN is a single point in the function space, while an RNN changes at each learning time (discrete in our definition) and so represents an orbit in the function space rather than a single point. Using the language of dynamical systems, we show that RNNs and their unfolded FNN counterparts belong to different mathematical categories. An FNN represents a fixed, time-T map of the RNN’s dynamics but cannot generate the full orbit that characterizes the behavior of the discrete-time RNN. This challenges the sufficiency of functional equivalence assumed by Doerig et al. (2019), because plastic RNNs cannot be reduced to static input-output mappings without losing these essential properties. Usher (2021) informally makes the case for this for standard RNNs in his initial work on the unfolding argument. This proof adds rigor and definitively demonstrates it to be the case for plastic RNNs.

**Figure 3:**
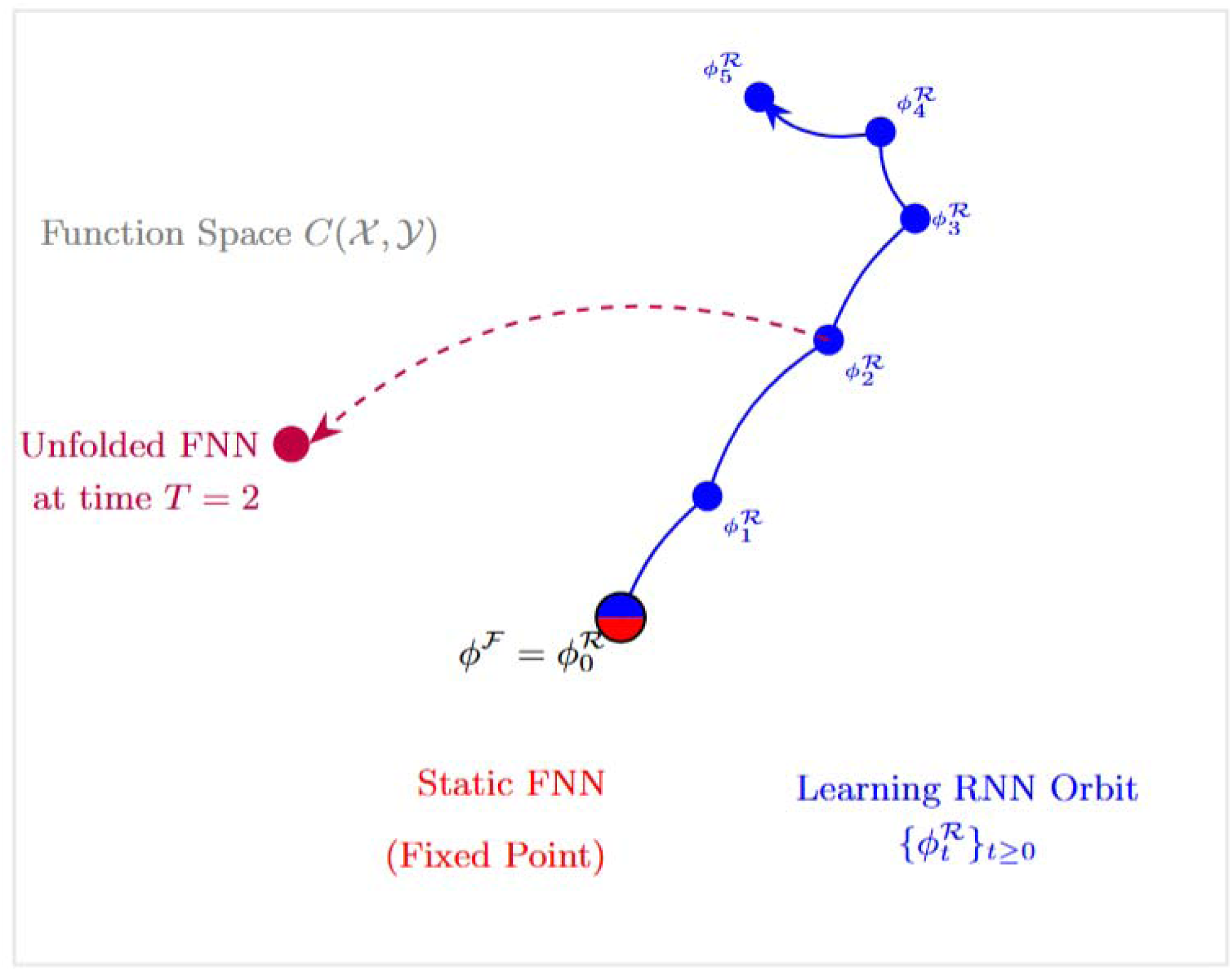
Visual representation of the argument from dynamical systems perspective. A static FNN represents a fixed point in function space (red), while a learning RNN generates an orbit (blue trajectory). The unfolding theorem can create an FNN equivalent to any single point on the RNN’s orbit (purple), but cannot capture the orbit-generating dynamics in function space.

Next, we leverage principles of feedback control and stability to target the assumption that an open-loop FNN can replicate closed-loop RNN dynamics for the case of discrete time networks (S1 Supplementary Appendix, Learning Implies Perturbation Response). We demonstrate that plastic RNNs and static FNNs respond fundamentally differently to temporal perturbations in their input-output behavior. While FNNs forget the effects of any input change after a fixed number of steps (determined by their input window size), RNNs maintain persistent traces of perturbations in their hidden states indefinitely (Figure 4). This temporal persistence property of RNNs - their ability to integrate and carry forward information from the distant past - cannot be replicated by static FNNs regardless of their size. This has direct relevance to consciousness, which arguably requires sustained integration of information across time rather than the bounded memory of windowed processing (Albantakis et al. 2023; Northoff & Zilio 2022; Hirschhorn et al. 2020; O’Reilly-Shah 2025b).

**Figure 4:**
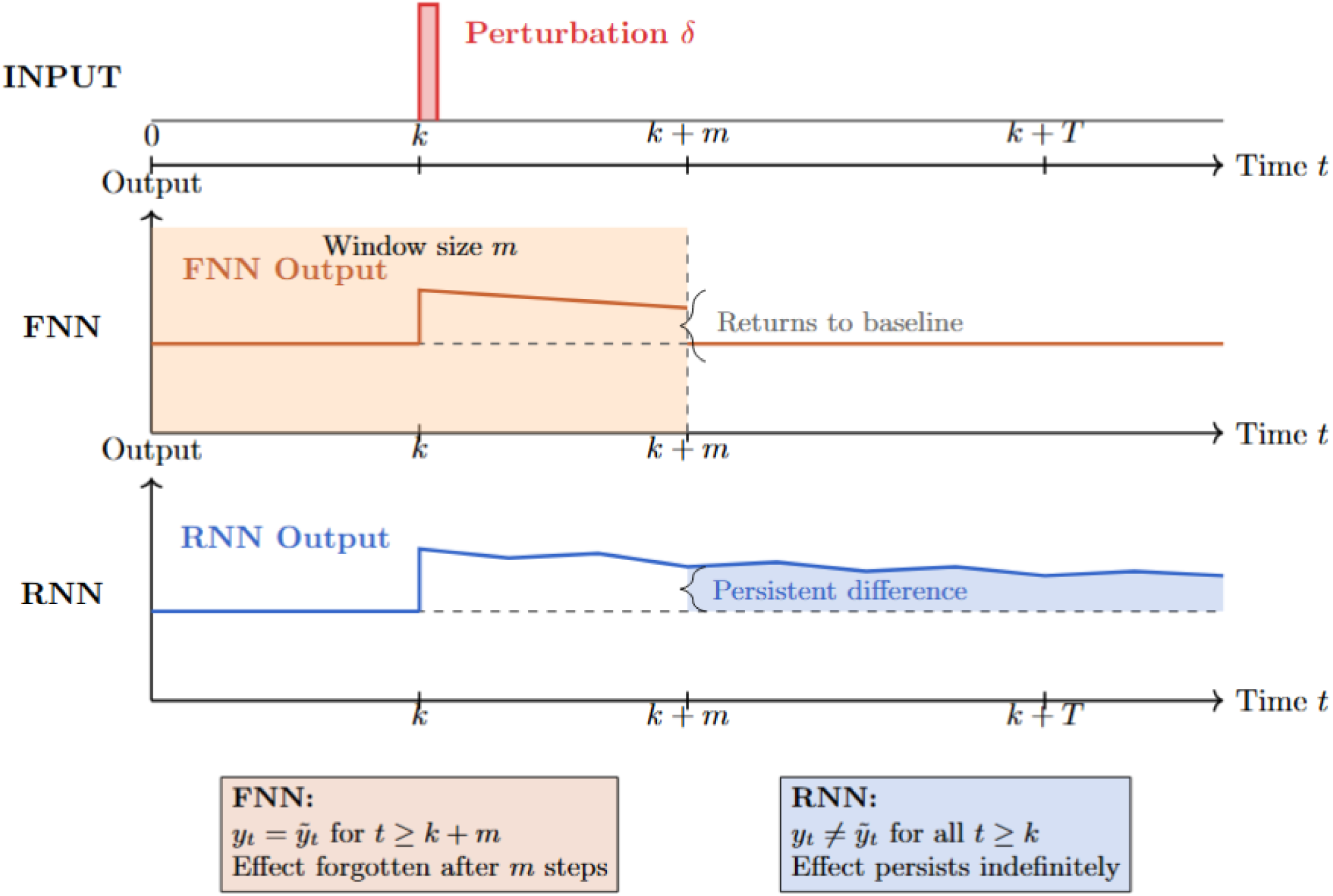
Visual representation of the argument from perturbational stability. A localized input perturbation δ is introduced at time *k* (red spike). The static Feedforward Neural Network (FNN, orange) shows a transient response that completely disappears after *m* steps (*t* ≥ *k*+*m*), due to its bounded sliding window memory. The Recurrent Neural Network (RNN, blue) maintains persistent traces of the perturbation indefinitely through its hidden state dynamics.

This challenges premise P3 by showing that evolving RNNs and static FNNs respond differently to perturbations in their input-output function behavior, potentially making them experimentally distinguishable through stability tests (Schurger et al. 2015; Eisen et al. 2024; Eisen et al. 2025) and other control-theoretic measures. Indeed, the clinical use of perturbational complexity index (PCI) (Casarotto et al. 2016) to assess for signatures of consciousness is reliant on a long-window response to perturbation (Casali et al. 2013), a feature present in RNNs with specific intrinsic organizational structure and absent in FNNs (Maschke et al. 2024; Casali et al. 2013). It should be noted that this does not violate P4 as we are specifically discussing differences in the input-output behavior, not differences in the internal workings of the networks.

This function space perturbation is more powerful than Usher’s (2021) state space perturbation point because it operates at the level of the input-output mapping itself rather than the internal dynamics. Usher (2021) correctly observes that a fixed RNN and its unfolded FNN counterpart traverse different state space trajectories - the hidden states diverge even when both systems compute the same function. However, a proponent of the UA could respond that these are precisely the kind of internal differences that P4 declares inadmissible: so long as the input-output function remains identical, the differing state trajectories are not experimentally accessible without examining the system’s internal workings. Our perturbation argument sidesteps this objection. When plasticity is present, input perturbations do not merely alter the hidden state trajectory while leaving the computed function intact; they alter the function itself. A plastic RNN modifies its weights during processing, changing what the system computes for all subsequent inputs. This change is observable at the input-output level without any appeal to internal measurement. The FNN, lacking this mechanism, returns to its fixed baseline after the input perturbation exits its input window. This divergence in observable input perturbation response is therefore immune to the P4 objection that constrains Usher’s original argument.

Our last area of formal investigation addresses the growth in complexity inherent to plastic systems and the implications of this for realistic, physically constrained biological systems. We provide a constructive proof demonstrating that plasticity within an RNN results in the ability of the network to approximate an arbitrarily large family of functions, whereas any single, static FNN is only able to approximate itself (S1 Supplementary Appendix, Learning Implies Approximation Capacity) (Song et al. 2023). This creates an unbridgeable gap between the functional repertoire of a plastic RNN and the capacity of any single, static FNN. A related result returns to the first argument, Corollary 3.5 (S1 Supplementary Appendix, Learning Limits Unfolding, No Finite Family of FNNs Can Track a Plastic RNN). When paired, these results form a powerful demonstration of the limits of FNNs in terms of their capability to reconstitute the functionality inherent to plastic RNNs (Figure 5). These results imply that even with perfect knowledge of the future behavior of a plastic RNN, it is impossible to switch to a finite library of FNNs to recapitulate that functionality. (NB: due to stochasticity and to chaotic behavior [yielding a deterministic yet non-predictable system] inherent to biological neural network systems, such perfect knowledge itself is impossible to attain.)

**Figure 5:**
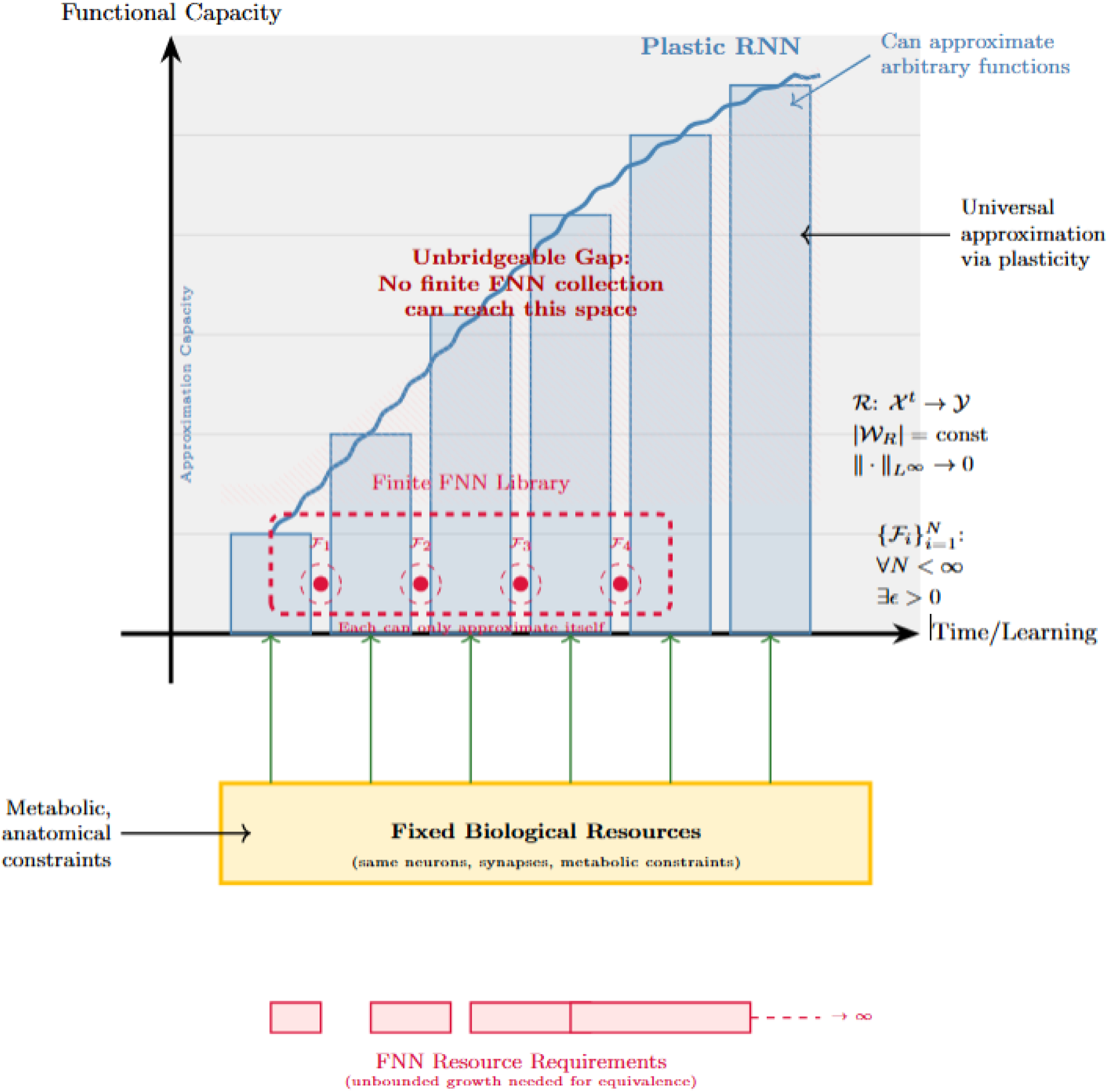
Visual representation of the argument from complexity growth and resource constraints. A plastic RNN (blue) achieves expanding functional capacity over time while operating with fixed biological resources (gold bar, constant width). The RNN’s approximation capacity grows vertically through learning, eventually capable of approximating arbitrary functions within the continuous function space *C*(*X*, *Y*). In contrast, static FNNs (red points) form a finite library {*F*, *F*, …, *F* } where each network can only approximate itself, creating a bounded functional repertoire. The hatched region represents the mathematical proof that no finite collection of static networks can access the functional space achievable by a plastic RNN with fixed resources. While the RNN maintains constant resource requirements, equivalent FNN implementations would require unbounded resource growth (approaching infinity).

While the concept of resource capacity is not well-defined in the neural network literature, these arguments also have implications for the recapitulation of biological RNNs in FNN systems. By ’resource capacity,’ we refer to the physical and computational constraints of a network, such as its total number of neurons and connections, as well as the metabolic or computational costs associated with its operation. A biological RNN operates using a physically constrained set of resources that, while dynamic and plastic, remains bounded over time. The functionality of this biological RNN cannot be replicated by a finite library of FNNs and, through learning, can approximate any arbitrary set of input-output functions. Thus, real, physically constrained, plastic biological RNNs cannot be re-instantiated in a FNN. The brain has a well-recognized capacity for lifelong (but probably bounded) learning within what might be considered quite strict metabolic, anatomical, and developmental constraints. These formal findings challenge the practical applicability and relevance of premise P2, as maintaining functional equivalence over time would require an unbounded library of FNNs.

#### The Turing Completeness Generalization

These plasticity arguments address the original formulation of the unfolding argument. However, Herzog et al. (2022) have proposed a stronger version, generalizing the UA beyond the RNN/FNN comparison to all Turing Complete (TC) systems. They argue that "any function, and even every process can be implemented with many different causal structures," concluding that "empirical results about consciousness are not provably due to causal structure."

On its face, this generalization would render our plasticity-based boundary conditions moot: one could concede that no single static FNN tracks a plastic RNN while maintaining that some TC system with different internal organizational structure could serve as a behaviorally identical substitute. We argue that this inference, while intuitive, is actually comprised of two distinct claims: that any computable function can be realized on diverse substrates (which Turing Completeness does guarantee) and that any specific realization of a computation can be reproduced with different internal organization in a manner that is indistinguishable under experimental probing (which it does not). Computational class membership guarantees the existence of some program computing a given function; it does not guarantee the existence of a realization that is process-equivalent to an arbitrary implementation in a different system.

This distinction is consistent with the multiple realizability literature, which likewise distinguishes between computational equivalence and causally relevant computational differences (Shapiro 2000; Kim 1992). Perturbation provides an empirical lever. As we demonstrate for the plasticity case (S1 Supplementary Appendix, Section 6), systems with different internal organization respond differently to mid-course interventions, and these differences manifest at the input-output level, not solely in internal observables. A system that reuses the same parameters at every step (an RNN) and one that instantiates independent parameter copies at each step (an unfolded FNN, or an alternative TC realization) will generically exhibit different perturbation profiles. This extends to any proposed TC substitute: matching the input-output function of a plastic RNN at a fixed time horizon does not entail matching its response to perturbation, and perturbation response is experimentally accessible without violating P4. We do not address the question of whether such perturbational differences are relevant to consciousness. Differences under perturbation establish that TC systems implementing the same computation are empirically distinguishable. A fuller formal treatment of this non-entailment is beyond the scope of the present manuscript and will be developed in forthcoming work.

Additionally, for systems engaged in ongoing environmental interaction (where future inputs depend on prior outputs and on unpredictable environmental dynamics) the input stream to the plastic RNN or any TC implementation of it cannot be pre-specified on a static tape. Any TC system that causally tracks the behavior of a plastic RNN in such a setting must itself receive the same environmental inputs and update its internal state accordingly. Such a system would, by construction, share the original’s dynamical internal organization and would not constitute the kind of "different causal structure, same function" substitute that the UA or TC generalization requires.

The Interactive Turing Machine (ITM) literature formalizes this distinction between function-computation and process-execution, referring to *a priori* unpredictable environmental inputs as ω-streams (Goldin & Wegner 2008; van Leeuwen & Wiedermann 2001; Cabessa & Siegelmann 2012; Cabessa & Siegelmann 2014). While some in the ITM community were interested in so-called super-Turing computation, modern formulations of ITMs do not appeal to hypercomputation (Fu 2022; Martin et al. 2022; Martin et al. 2023; Jaeger et al. 2023). The implications of environmental coupling on computability and computational multiple realizability are important but, again, beyond the scope of this manuscript. A fuller discussion of ITMs and these implications will be developed in forthcoming work. We note here only that this environmental-coupling process-based consideration may provide an obstruction to unfolding and the TC generalization that is independent of, and orthogonal to, the plasticity arguments we have presented.

#### Lenient Dependency and the Restoration of Empirical Testability

Our results have bearing on a critical result in the consciousness literature: the Kleiner and Hoel (2021) substitutability argument. This argument demonstrates that if behavioral reports and internal observables are independent, then any minimally informative theory is already falsified (K&H Theorem 3.10) and if strictly dependent, then the theories are unfalsifiable (K&H Theorem 4.3). Only lenient dependency, where reports constrain but do not determine internal observables, permits empirical testability. Kleiner and Hoel identify no current theory meeting this criterion.

However, plastic systems exhibit this intermediate relationship. The intuition is straightforward: the longer you observe a plastic system’s behavior, the more you can ’rule out’ possible kinds of systems that might be producing it. A static surrogate might match the plastic system’s outputs for a short window, but it cannot match the persistent perturbation traces and growing information capacity that our formal results establish. As the observation horizon lengthens, systems that lack plasticity produce observably different behavioral profiles, and the space of candidate systems consistent with the full behavioral record narrows.

More formally, our perturbation and information-theoretic results provide the concrete exclusion mechanism. A localized perturbation at time *k* induces output differences at all subsequent times in a plastic RNN, whereas a static FNN forgets the perturbation after a fixed number of steps determined by its window size *m* (S1 Supplementary Appendix, Section 6). For any observation horizon longer than this window, the static surrogate’s behavior diverges from the plastic system’s: it can no longer account for the observed behavior. Similarly, the mutual information between target and RNN output grows with observation horizon, while a window-limited FNN hits a ceiling imposed by its fixed window (S1 Supplementary Appendix, Section 3). Together, these results progressively exclude static surrogates as the observation horizon lengthens.

This rules out the first horn of the Kleiner-Hoel dilemma: extended behavioral observation of a plastic system excludes static surrogates, because no static unfolding or finite family of feedforward networks can reproduce the behavioral profile over time (S1 Supplementary Appendix, Section 3). But the second horn, that behavior uniquely determines internal organization, is not implied either. Even systems that obey the same plasticity laws and produce identical behavioral traces may differ internally in ways that leave no observable signature, such as how individual units are labeled or how internal states are coordinated. Behavior constrains what the underlying system could be, without pinning it down to exactly one possibility. This is the intermediate relationship Kleiner and Hoel termed lenient dependency: the only regime in which theories of consciousness remain empirically testable. Hoel (2025), citing a preprint version of this paper, recently arrived at a convergent conclusion via a different mathematical approach.

NB: This escape route from the unfolding and substitutability arguments does not yield any additional *a priori* credibility for any theory of consciousness. Theories incorporating plasticity simply avoid the UA’s critique and remain viable candidates for empirical investigation. This is a restoration of testability, not a claim of correctness.

## Discussion

Our formal analysis maps the scope conditions of the unfolding argument, identifying where its premises hold and where they require qualification. We build upon Doerig et al.’s (2019) important contribution by identifying specific boundary conditions; specifically, the analyses presented establish a natural division in the theoretical landscape. It remains true that any theory claiming special computational significance of recurrence *per se* fails to meet the challenge of the unfolding argument or the Turing Completeness generalization. No theory gains credibility on our analysis; we metaphorically open the door, but do not walk through it. Our analysis says simply that empirical testability is restored for theories that explicitly incorporate plasticity.

### Specific Implications for Major Theories of Consciousness

This finding has significant implications for physicalist theories of consciousness. It restores to testability theoretical frameworks where the *how* of information processing - involving time, dynamics, and fast plasticity - is as important as the *what* of the input-output function. Our findings do not challenge the view that, computationally, the plastic RNN structure could be realized in different causal forms. The arguments we have made here have no implications for the specific *implementation* of the computations performed by a plastic RNN.

The arguments related to resource constraints echo Dennett (1993): "sometimes an *impossibility in fact* is theoretically more interesting than a *possibility in principle*." Imagine the brain’s recurrent neural networks, which may be represented as a vast set (in the mathematical sense) of recurrent neural networks. (It is due to plasticity that we should represent it as a set of RNNs rather than as a single RNN; plasticity means that the underlying weights, connections, and even node count of the network will change over time.) While it may be possible to unfold each RNN in this set into an equivalent finite set of feedforward architectures, this would still fail to capture open-ended, plastic processes. Each static FNN would represent an instant, but not the evolving state of the brain. Static FNNs lack the capacity for this continual state integration and adaptation.

For Integrated Information Theory (IIT) (Albantakis et al. 2023), the original challenge from Doerig and colleagues (2019) contended that a system with a high degree of integrated information (Φ) could be functionally mimicked by a feedforward system with negligible Φ, making IIT either false or unscientific. A relevant distinction is the *capacity* for consciousness versus the *actual instantiation* of conscious states (Mudrik et al. 2023). Measures like Φ in IIT are computed from the system’s causal structure - specifically, the transition probability matrix (TPM) - and purport to quantify the degree of consciousness a system possesses by virtue of its architecture. According to IIT, Φ is meant to measure both dispositional and occurrent properties of conscious systems: the amount of integrated information is the amount of consciousness actually present at that moment. So if a network happens to be in a highly integrated state, IIT would say it is experiencing a correspondingly high level of consciousness. By contrast, measures like PCI capture only an occurrent property: what the system is currently doing, not what it is capable of (Casarotto et al. 2016; Casali et al. 2013). .

Computationally, the underlying causal structure of a candidate system and its TPM can still be expressed in any unfolded FNN or Turing Complete system. Consequently, IIT remains challenged and our findings do not alter the conclusions of Doerig et al. (2021; 2019) (Note, the foregoing is based on a strong interpretation of IIT; see e.g. Mediano et al. (2022) for a more nuanced discussion of the point. IIT proponents themselves have used the unfolding equivalence didactically, emphasizing that ’being is not doing.’ Our analysis suggests that incorporating plasticity into IIT models could resolve this tension, though IIT proponents would need a principled rather than ad hoc reason for doing so.)

Global Neuronal Workspace Theory (GNWT) (Mashour et al. 2020; Dehaene & Naccache 2001) proposes that consciousness involves information being "globally broadcast" within a dynamic workspace involving recurrent processing. Our analysis adds nuance to the claim in Doerig et al. that it is protected by being a functional, rather than structural, theory. Thinking mechanistically, a static implementation of a global workspace would be vulnerable to the unfolding argument. However, a dynamic workspace that incorporates plasticity, allowing for learning-dependent changes in information access patterns, is testable in our view. This framing makes the *evolution* of the workspace during perceptual information processing episodes the key to developing testable predictions.

Recurrent Processing Theory (RPT), developed by Lamme (2018), was also one of the original targets of the unfolding argument. Our results do not rescue RPT from the unfolding argument. RPT posits that consciousness arises when recurrent interactions between visual areas integrate features into coherent perceptual wholes, distinguishing it from unconscious feedforward processing (Lamme 2018). In fairness, Lamme (2018) explicitly identifies plasticity as a potential "missing ingredient" that distinguishes conscious from unconscious recurrent processing, but we leave it to the defenders of RPT to develop the point.

The state space theory (SST) of consciousness (O’Reilly-Shah 2025b; O’Reilly-Shah 2026) posits that consciousness arises from delay coordinate embedding (DCE) in plastic, recurrent neural networks. Recent mathematical (Duan et al. 2023; Hart 2025) and empirical (Uribarri & Mindlin 2022; Ostrow et al. 2024) work demonstrate that RNNs (LSTMs, reservoir computers) trained on dynamical time series data make use of embedding structure for prediction. These findings potentially link the behavior of neural RNNs to perception (O’Reilly-Shah & Selvitella 2026). DCE inherently requires temporal depth because it is incoherent to speak of DCE without time series data. Plasticity ensures the embedding evolves to match internal and environmental dynamics monitored and reported on via neuronal spike trains, creating an exploitable relation (Shea 2018). These are fundamental commitments and arise directly from the core hypothesis (O’Reilly-Shah 2025a). This caveat paper formalizes obstructions mentioned in the original article where SST was developed (O’Reilly-Shah 2025b).

#### Limitations, Extensions, and Open Questions

A significant limitation concerns the relationship between the timescales of consciousness and plasticity. Many conscious processes occur rapidly, while significant structural plasticity can take much longer. This raises the empirical questions of whether rapid forms of synaptic plasticity are sufficient to provide protection from unfolding and whether conscious processes depend more on ongoing plastic changes or their accumulated historical effects. As advocated for by He (2023), there may be multiple different sorts of neurobiological mechanisms working in concert or separately in support of consciousness. Short-term synaptic plasticity and spike timing dependent plasticity have been well documented to occur on relevant timescales for perception and consciousness, including one review identifying these as relevant for sensory timing (Motanis et al. 2018; Regehr 2012; Barri & Mongillo 2022; Li et al. 2024).

Our mathematical proofs establish boundaries in principle, but biological realities such as metabolic energy costs and stability-plasticity tradeoffs may impose additional constraints. As stated, our proofs are theory-neutral; they demonstrate which dynamic properties are lost in unfolding but do not, by themselves, establish which of these properties are necessary for consciousness.

One might object that even if plastic RNNs cannot be tracked by finite FNN libraries, one could instantiate the same computation by continuously replacing the RNN with its FNN, or just implementing it in another Turing Complete system. This alternative implementation would need to, at each moment, track precisely the dynamical trajectory and plasticity-mediated changes of the original system. Such a hypothetical alternative-implementation/FNN zombie with the dynamical and plasticity properties of the original RNN-based system would be identical in terms of empirical evidence for consciousness. However, a few observations:

1. Such a system is no longer a static FNN in any sense relevant to the unfolding argument. It reflects the computational function space trajectory of the original plastic RNN implemented in an alternative substrate. Standard Universal Approximation Theorems guarantee that a static FNN can approximate a specific target function to arbitrary accuracy (see e.g. Augustine 2024). They do not, however, guarantee that a single static architecture can approximate an evolving family of functions (a trajectory through function space) without itself undergoing parameter modification.
2. A static FNN could not change structure/weight in response to input without an external agent acting on the FNN (or would require instant-to-instant replacement.) A plastic RNN, or its implementation on a Turing Complete system, has the power to modify itself.

Whether consciousness requires dynamical properties (temporal thickness, real-time state-space and function-space traversal, history-dependent trajectories) or whether these are merely contingent features of biological consciousness remains an open empirical question. Our arguments support *only* the conclusion that plasticity provides a way to make the recurrence hypothesis testable and within the realm of science.

Our formal arguments are based on discrete-time RNNs. We remain agnostic on whether *continuous*-time dynamics are essential to consciousness. Some have argued that this induces a requirement for super Turing computation (Lucas 1961; Penrose 1989) or relates to the importance of biological substrate in consciousness (e.g., Seth 2025; Milinkovic & Aru 2025). Unlike substrate dependence arguments, our result does not privilege biological implementation; it identifies a formal property, plasticity, that is substrate-neutral. We are also agnostic on the question of instantiation versus simulation: whether a perfect simulation of a plastic RNN *is* conscious or *acts as if* conscious. This is a question our formal arguments cannot resolve. What we have established is more modest but, we believe, significant: theories with specific computational properties are empirically testable.

### Meta-Theoretical Implications and the Path Forward

By delineating these boundaries, our analysis shifts the burden of proof in consciousness science. Proponents of static theories should consider whether the boundary conditions identified here bear on their formulations. Conversely, proponents of dynamic theories must specify precisely which of these properties they deem necessary and provide empirical evidence to support their claims. This suggests new research directions focusing on the relationship between consciousness and learning, development, and pathological plasticity (Cleeremans 2007).

Our formal results engage directly with the critical tradition of multiple realizability. On Shapiro’s analysis, the question is whether the structural difference between the plastic RNN (whose weights and arity evolve) and static FNN (whose weights and arity are fixed) is relevant to the function at issue (Shapiro 2000). As we’ve established, it is indeed a relevant question for plastic systems. Dynamical internal organization constitutes a relevant property in precisely Shapiro’s sense: it determines the system’s future input-output behavior. The distinction between function-level and process-level MR (Shapiro 2000; Polger & Shapiro 2016) thus becomes a diagnostic criterion: theories that can be reduced to static functional descriptions inherit all the vulnerabilities of classical MR, while theories that invoke genuine temporal dynamics escape them.

Our analysis identifies which theoretical frameworks remain viable candidates for empirical investigation in light of the unfolding argument’s scope conditions. A theory constructed to be immune to the unfolding argument does not make it a stronger contender as a theory of consciousness. The unfolding argument and its caveats are a valuable guide. It helps identify which theoretical approaches can yield genuine insight into the nature of consciousness while remaining grounded in empirical science.

## Conclusions

In conclusion, our formal analysis of the unfolding argument’s boundaries tells us that theories under which consciousness depends on recurrence might not be invalidated by the unfolding argument if they incorporate, or at least allow for, fast plasticity in recurrent networks on the timescale of conscious neural processes. We emphasize that nothing in our analysis provides independent reason to believe plasticity is relevant to consciousness; our results identify boundary conditions on a logical argument, not evidence for any particular theory. The dynamic view is both scientifically respectable, grounded in rigorous mathematical proofs, and empirically productive, suggesting specific experimental approaches for advancing our understanding of consciousness.

Disclosure of funding

This research received no specific grant from any funding agency in the public, commercial, or not-for-profit sectors.

## Author contributions

All authors meet ICMJE criteria for authorship. Conceptualization: V-OS, A-Sch; Formal Analysis: A-Sel; Writing - Original Draft: : All authors; Writing - Review & Editing: All authors.

## Acknowledgements

The authors would like to thank their respective universities for their generous support of the authors’ time.

## Declaration of generative AI and AI-assisted technologies in the writing process

During the preparation of this work the author(s) used OpenAI ChatGPT, Anthropic Claude, and Google Gemini models in order to polish written content, assist with the formatting of proofs in LaTeX, and assist with the coding of TikZ figures in LaTeX. After using this tool/service, the authors reviewed and edited the content. The authors take full responsibility for the manuscript in its entirety. Citations were manually retrieved from traditional sources (e.g., PubMed).

## Data Availability Statement

There is no new data associated with the article.

