## Supplementary material for "A Caveat Regarding the Unfolding Argument: Implications of Plasticity": S1 Supplementary Appendix

### A Caveat Regarding the Unfolding Argument: Implications for Computational Theories of Consciousness

### Contents


**Roadmap** 3

**I Introduction** 5

**1 Notations and Preliminaries** 5

**2 The Unfolding Argument** 6

**A Note on Terminology: Learning and Plasticity** 7

**II Proofs** 9

**3 Argument: Learning Limits Unfolding** 9

**4 Argument: Learning Implies More Information** 13

**5 Argument: Learning at Discrete Times Needs Countable Unfolding** 16

**6 Argument: Learning Implies Perturbation Response** 19

**7 Argument: Learning Implies Approximation Capacity** 23

**8 Kleiner–Hoel and Lenient Dependency** 27

### Roadmap

This supplementary appendix provides the full mathematical foundations for the claims made in the main text. We offer this roadmap to orient the reader.

#### Overview

The unfolding argument (UA), as stated by its proponents, asserts that any recurrent neural network (RNN) can be "unfolded" into a feedforward neural network (FNN) computing the same input–output function. The basic mathematical claim is correct for fixed parameters and a fixed finite time horizon $T$ (Theorem 2.1). However, this equivalence is far more limited than is sometimes appreciated. In this appendix we develop six independent lines of argument showing that the UA cannot be extended in the ways required to draw the broad conclusions about consciousness that have been attributed to it, together with two additional sections addressing the computability-theoretic and empirical-testability implications.

#### Part I: Introduction and Preliminaries

We begin by fixing notation for feedforward and recurrent neural networks (Definitions 1.1 and 1.2), stating and proving the standard unfolding theorem (Theorem 2.1), and clarifying the distinction between *plasticity* (any parameter change over time) and *learning* (task-directed parameter optimization). Our formal results require only plasticity, not learning per se.

#### Part II: Proofs

The core of the appendix consists of six arguments, each attacking the generalizability of the UA from a different mathematical angle. While logically independent, they are mutually reinforcing: taken together, they show that the limitations of unfolding are not an artifact of any single formalization but reflect a deep structural asymmetry between static and dynamic computation.

**Argument 1: Learning Limits Unfolding (§3).** We show that a plastic RNN whose input domain grows over time cannot be equivalent to any single static FNN for all future times (Theorem 3.2). The proof is set-theoretic: the RNN's input space $(\mathcal{X}_R)^t$ strictly grows with $t$, while any FNN has a fixed input space. A corollary (Corollary 3.5) extends this to show that no *finite family* of FNNs can track a plastic RNN, by a pigeonhole argument. This establishes that unfolding is necessarily *local in time*.

**Argument 2: Learning Implies More Information (§4).** Using the data processing inequality and the conditional Markov property of RNN hidden states (Theorem 4.3), we prove that a learning RNN processing a full input history can encode strictly more mutual information about a target variable than a static FNN restricted to a fixed-length window

(Theorem 4.5, Corollary 4.7). This information-theoretic gap is a direct consequence of the RNN's ability to accumulate temporal context through learned recurrence.

**Argument 3: Learning at Discrete Times Needs Countable Unfolding (§5).** We recast the distinction between FNNs and RNNs in the language of discrete dynamical systems. A static FNN corresponds to a *fixed point* of the identity map on function space (Theorem 5.2, Corollary 5.4), while a plastic RNN traces a countable *orbit* through function space (Proposition 5.6). We show that a static FNN is a better approximator of itself than any learning RNN whose orbit never passes through the FNN's input–output map (Proposition 5.9), but this self-approximation property is precisely what makes static networks unable to adapt.

**Argument 4: Learning Implies Perturbation Response (§6).** We prove that localized input perturbations are *forgotten* by static FNNs after a fixed number of steps equal to their window size (Theorem 6.1), whereas in RNNs such perturbations generically persist indefinitely in the hidden state and output (Theorem 6.2). This persistence survives unfolding (Corollary 6.3) and provides an experimentally testable signature distinguishing recurrent from feedforward architectures—directly relevant to theories of consciousness that require temporal integration.

**Argument 5: Learning Implies Approximation Capacity (§7).** We show that a static FNN can approximate only itself in the $L^\infty$ norm (Theorem 7.2), and that any two disjoint finite families of continuous functions are positively separated (Theorem 7.5). By contrast, learning RNNs are universal approximators whose approximation capacity grows without bound. Unfolding does not confer adaptation: the unfolded network inherits only the behaviors present at construction.

**Lenient Dependency (§8).** We connect our results to the empirical testability framework of Kleiner and Hoel, showing that our plasticity results exhibit the structure of *lenient dependency*: extended behavioral data constrains admissible internal organizations (ruling out static surrogates) without uniquely determining them (Lemma 8.1, Remark 8.3).

### Part I
### Introduction

#### 1 Notations and Preliminaries

For completeness, we provide here the definitions of Feedforward Neural Networks and Recurrent Neural Networks that we will consider throughout our analysis.

**Definition 1.1** (Feedforward Neural Network)
*Let $\mathcal{X}_F \subseteq \mathbb{R}^m$ be the input space, $\mathcal{Y}_F \subseteq \mathbb{R}^n$ the output space, and $L \in \mathbb{N}$ the number of layers. A* feedforward neural network (FNN) *of depth $L$ is a tuple*

$$\mathcal{F} = (\{\mathcal{H}_\ell\}_{\ell=1}^L, \{W_{F,\ell}\}_{\ell=1}^L, \{f_\ell\}_{\ell=1}^L)$$

*where:*

- *$W_{F,\ell} \in \mathcal{W}_F$ are the parameters for every $l = 1, \ldots, L$, with $\mathcal{W}_F$ representing the parameter space.*
- *$\mathcal{H}_\ell := \mathbb{R}^{d_\ell}$ is the representation space at layer $\ell$, with $\mathcal{H}_0 = \mathcal{X}_F$ and $\mathcal{H}_L = \mathcal{Y}_F$.*
- *For each $\ell = 1, \ldots, L$: $f_\ell : \mathbb{R}^{d_{\ell-1}} \times W_{F,\ell} \to \mathbb{R}^{d_\ell}$ is the $\ell$-th layer function, with $f_L$ the output layer and with $d_0 = m$ and $d_L = n$.*

*Given $x_F \in \mathcal{X}_F$, the FNN computes*

$$\begin{aligned} h^{(0)}(x_F) &:= x_F, \\ h^{(\ell)}(x_F) &:= f_\ell(h^{(\ell-1)}(x_F), W_{F,\ell}), \quad \ell = 1, \ldots, L, \\ y_F &:= h^{(L)}(x_F), \end{aligned}$$

*with $y_F \in \mathcal{Y}_F$. The FNN's input-output map is $\phi^{\mathcal{F}_L} : \mathcal{X}_{F_L} \to \mathcal{Y}_F$, with $\phi^{\mathcal{F}_L}(x) = h^{(L)}(x)$.*

**Definition 1.2** (Recurrent Neural Network)
*Let $\mathcal{X}_R \subseteq \mathbb{R}^m$ be the input space, $\mathcal{Y}_R \subseteq \mathbb{R}^n$ the output space, and $\mathcal{H}_R \subseteq \mathbb{R}^d$ the hidden state space. A* recurrent neural network (RNN) *is a tuple*

$$\mathcal{R} = (\mathcal{H}_R, \mathcal{W}_R, h_0, W_R, f, g)$$

*where:*

- *$W_R \in \mathcal{W}_R$ are the parameters, with $\mathcal{W}_R$ representing the parameter space.*
- *$h_0 \in \mathcal{H}_R$ is the initial hidden state.*
- *$f : \mathcal{H}_R \times \mathcal{X}_R \times \mathcal{W}_R \to \mathcal{H}_R$ is the state update function.*

- $g : \mathcal{H}_R \times \mathcal{X}_R \times \mathcal{W}_R \to \mathcal{Y}_R$ *is the output function.*

*Given an input sequence* $(x_1, \ldots, x_T)$ *with* $x_t \in \mathcal{X}_R$ *for every* $t \in \mathbb{N} : 1 \le t \le T < +\infty$*, the RNN computes, for* $t = 1, \ldots, T$*,*

$$h_t = f(h_{t-1}, x_t, W_R),$$
$$y_t = g(h_t, x_t, W_R),$$

*with* $y_t \in \mathcal{Y}_R$*. The RNN's input-output map at time* $t$ *is the function*

$$\phi_t^{\mathcal{R}} : \mathcal{X}^t \to \mathcal{Y}$$

*such that*

$$\phi_t^{\mathcal{R}}(x_1, \ldots, x_t) := y_t = g(h_t, x_t, W_R).$$

### 2 The Unfolding Argument

The unfolding argument states that any RNN with fixed parameters and finite time horizon $T$ can be unfolded into a FNN computing the same input-output function over input sequences of length $T$.

**Theorem 2.1** (Unfolding a Recurrent Neural Network into a Feedforward Neural Network) *Let* $\mathcal{R} = (\mathcal{H}_R, \mathcal{W}_R, h_0, W_R, f, g)$ *be a RNN, with input space* $\mathcal{X}_R \subseteq \mathbb{R}^m$ *and output space* $\mathcal{Y}_R \subseteq \mathbb{R}^n$*. For any time horizon* $T \in \mathbb{N}$*, there exists a feedforward neural network* $\mathcal{F}_T$ *whose input space is* $(\mathcal{X}_R)^T$ *and output space* $\mathcal{Y}_R$*, such that the input-output function*

$$\phi^{\mathcal{F}_T} : (\mathcal{X}_R)^T \to \mathcal{Y}_R$$

*satisfies*

$$\phi^{\mathcal{F}_T}(x_1, \ldots, x_T) = g(h_T, x_T, W_R)$$

*where the hidden states are defined recursively by*

$$h_t = f(h_{t-1}, x_t, W_R), \qquad h_0 \text{ given}, \quad t = 1, \ldots, T.$$

*Proof.* We explicitly construct the FNN $\mathcal{F}_T$ as follows. Let the input space be $(\mathcal{X}_R)^T$, i.e., $T$ concatenated inputs $x_1, ..., x_T$. For each $t = 1, \ldots, T$, define the hidden layer computation as

$$h_t := f(h_{t-1}, x_t, W_R), \quad h_0 \text{ fixed}.$$

The final output is computed as $g(h_T, x_T, W_R) \in \mathcal{Y}_R$. This composition can be implemented as a FNN of depth $L = T$, using shared parameters $W_R$, with each layer corresponding to a timestep's update, followed by the output map $g$. The resulting input-output function is exactly

$$\phi^{\mathcal{F}_T}(x_1, ..., x_T) = g(h_T, x_T, W_R),$$

which is, by construction, the same as the RNN output after $T$ steps given the sequence $(x_1, ..., x_T)$. The function spaces match by construction: $\phi^{\mathcal{F}_T} : (\mathcal{X}_R)^T \to \mathcal{Y}_R$.

■

**Corollary 2.2** (Unfolding with Uniform Hidden Size)
*Let $\mathcal{R} = (\mathcal{H}_R, \mathcal{W}_R, h_0, W_R, f, g)$ be a RNN with hidden state space $\mathcal{H}_R \subseteq \mathbb{R}^d$ of fixed dimension $d$ for all time steps. Then, for any $T \in \mathbb{N}$, the unfolded FNN $\mathcal{F}_T$ can be realized so that $W_F = W_R \times W_R \times \cdots \times W_R$, with $dim(W) = d$.*

*Proof.* This is basically a tautology and follows from the construction in the proof of Theorem 2.1.

■

**Remark 2.3** (Unfolding with Larger Layers and Deactivated Neurons)
*Alternatively, for any $D \geq d$, the unfolded FNN can be constructed so that each hidden layer has $D$ neurons, where only the first $d$ neurons are actively used and the remaining $D - d$ neurons are deactivated (e.g., set always to zero or ignored by subsequent connections). This construction is functionally equivalent to the one above, since the inactive units do not affect computation (their outgoing weights can be set to zero). Potentially, layer $l$ can have a total of $D_l$ neurons with only $d$ active and $D_l - d$ inactive for every $l = 1, \ldots, L$ and not all $D_l$'s equal. This flexibility may be desirable in practice for architectural compatibility or implementation convenience.*

**Remark 2.4** (Necessity of Sufficient Layer Width in Unfolding)
*In general, an exact unfolding of an RNN with hidden state dimension $d$ into a FNN over $T$ steps is* not *possible if the FNN uses fewer than $d$ neurons per hidden layer. Each unfolded layer must be able to represent all possible RNN hidden states at that time; reducing the layer width would constrain the function class and make it, in general, impossible to simulate the original RNN's dynamics and output over arbitrary input sequences.*

### A Note on Terminology: Learning and Plasticity

Throughout this appendix, we use the terms "learning" and "plasticity" in related but distinct ways. We clarify their usage here to avoid confusion.

**Definition 2.5** (Plasticity)
Plasticity *refers to any process by which the parameters (weights) of a neural network change over time. Formally, if $W_t$ denotes the network parameters at time $t$, plasticity is present whenever there exists $t_1 < t_2$ such that $W_{t_1} \neq W_{t_2}$.*

**Definition 2.6** (Learning)
Learning *refers to plasticity that is directed toward a task or objective, typically formalized as optimization of a loss function. Learning is thus a specific form of plasticity, but not all plasticity constitutes learning in this sense.*

**Remark 2.7** (Scope of Our Formal Arguments)
*The formal arguments presented in Part II require only* plasticity*—that is, non-trivial change in network parameters over time—and do not require that this change be task-directed learning. While we sometimes use "learning" in section titles and exposition for readability (e.g., "Learning Limits Unfolding"), the mathematical results depend only on the weaker condition that $W_t$ evolves non-trivially over time.*

*This distinction is important for two reasons:*

1. ***Biological relevance***: *Neural plasticity encompasses many mechanisms beyond task-directed learning, including short-term synaptic facilitation, spike-timing-dependent plasticity (STDP), and homeostatic regulation. Our arguments apply to all such mechanisms operating on timescales relevant to perception and consciousness.*

2. ***Relationship to prior work***: *Usher (2021) demonstrated that even fixed-weight RNNs exhibit perturbational differences from their unfolded FNN equivalents due to transient dynamics in* state space*. Our contribution extends this by showing that plastic RNNs generate trajectories in* function space*—the space of input-output mappings itself changes over time. This is a stronger result: while Usher's state-space trajectories occur within a fixed computational regime, our function-space trajectories represent evolution of what computation the system performs.*

*In summary, where our proofs reference "learning," the reader may substitute "plasticity" without loss of validity. The critical requirement is temporal evolution of parameters, not the presence of an objective function or error signal.*

### Part II
### Proofs

#### 3 Argument: Learning Limits Unfolding

In this section, we prove that the unfolding argument is reversible if an only if there is no learning. Different networks learn differently when they are allowed to learn indefinitely. In particular, we stress the argument that a static FNN can be equivalent to an RNN at most in a point in time during learning. This implies that the unfolding argument would require a different FNN at each learning time, until the next learning time, when the network changes again. The argument shows that, under certain assumptions on the learning rule, there isn't a single static network that can approximate all the epochs of a network that can learn indefinitely, limiting the unfolding to static networks. We start with the definition of functional equivalence between networks.

**Definition 3.1** (Functional Equivalence with Domain Matching)
*Let $\mathcal{F}$ be an FNN with input-output map $\phi^{\mathcal{F}} : \mathcal{X}_{\mathcal{F}} \to \mathcal{Y}$, and $\mathcal{R}$ an RNN with time-t input-output map $\phi_t^{\mathcal{R}} : \mathcal{X}_{\mathcal{R}}^{(t)} \to \mathcal{Y}$. Suppose $t, s \in \mathbb{N}$ with $t \geq s$. We say that $\mathcal{F}$ and $\mathcal{R}$ are **Equivalent at** $t$, denoted $\mathcal{R} \equiv_t \mathcal{F}$, if and only if:*

1. *The input spaces coincide at time $t$: $\mathcal{X}_{\mathcal{F}} = \mathcal{X}_{\mathcal{R}}^{(t)}$,*
2. *The input-output functions agree: $\phi_t^{\mathcal{R}} = \phi^{\mathcal{F}}$ as functions $\mathcal{X}_{\mathcal{F}} \to \mathcal{Y}$.*

*We say that $\mathcal{F}$ and $\mathcal{R}$ are **Equivalent on** $[s, t]$, denoted $\mathcal{R} \equiv_{[s,t]} \mathcal{F}$, if and only if $\mathcal{R} \equiv_\tau \mathcal{F}$ for every $\tau \in [s, t]$.*

**Theorem 3.2** (Plasticity Precludes Static Equivalence)
*Let $\mathcal{R}$ be an RNN with $\phi_t^{\mathcal{R}} : \mathcal{X}^t \to \mathcal{Y}$ representing the RNN's input-output map at time $t$. Suppose that for every $T \in \mathbb{N}$ such that $T > 0$, there exists $t_1, t_2 \in \mathbb{N}$ with $t_1, t_2 > T$, such that $(\mathcal{X}_{\mathcal{R}})^{t_1} \subsetneq (\mathcal{X}_{\mathcal{R}})^{t_2}$. Then for any FNN $\mathcal{F}$, there is no $t^* \geq 0$ such that $\mathcal{R} \equiv_{[t^*,\infty)} \mathcal{F}$.*

*Proof.* Suppose to the contrary that there exists $t^* \geq 0$ and a static FNN $\mathcal{F}$ such that for all $t \geq t^*$, $\mathcal{R}$ is functionally equivalent to $\mathcal{F}$ on the sequence set $\mathcal{S}$, i.e., $\mathcal{R} \equiv_{[t^*,\infty)} \mathcal{F}$. For functional equivalence to hold for all $t \geq t^*$, the input domains must coincide, that is, the input domain of $\mathcal{F}$ must match the input domain of $\mathcal{R}$ at every such $t$. Since the FNN $\mathcal{F}$ has a fixed input space, say $\mathcal{X}_{\mathcal{F}}$, we must have

$$\mathcal{X}_{\mathcal{F}} = (\mathcal{X}_{\mathcal{R}})^t \quad \text{for all } t \geq t^*.$$

However, by the assumption of the theorem, for every $T > 0$, and in particular, for $T = t^*$, there exist $t_1, t_2 > T = t^*$ with $t_1 < t_2$ such that

$$(\mathcal{X}_{\mathcal{R}})^{t_1} \subsetneq (\mathcal{X}_{\mathcal{R}})^{t_2}.$$

But this means $(\mathcal{X}_{\mathcal{R}})^{t_1} \neq (\mathcal{X}_{\mathcal{R}})^{t_2}$. Hence, for any FNN $\mathcal{F}$, there does not exist $t^* \geq 0$ such that $\mathcal{R} \equiv_{[t^*,\infty)} \mathcal{F}$ on the set of sequences $\mathcal{S}$. ∎

**Remark 3.3**
*The assumption that for every* $T \in \mathbb{N}$ *such that* $T > 0$*, there exists* $t_1, t_2 \in \mathbb{N}$ *with* $t_1, t_2 > T$*, such that* $(\mathcal{X}_{\mathcal{R}})^{t_1} \subsetneq (\mathcal{X}_{\mathcal{R}})^{t_2}$ *is a low regularity version of the requirement that the RNN undergoes non-trivial learning for unbounded time intervals. The plasticity intrinsic in RNN during learning is so fundamental that it starts from the fact that the input-ouput function undergoes a change in the input space during learning. This fact is set-theoretical and independent of regularity assumptions on the activation functions. Obviously, this is distinct from a static FNN, which has fixed architecture, but also a type of plasticity different from non-recurrent FNNs, whose learning is based on parameter adaptation only, and not on input space adaptation.*

**Remark 3.4**
*This proof implies that unfolding is local in time. The same FNN* $\mathcal{F}$ *is equivalent to the corresponding RNN* $\mathcal{R}$ *and the unfolding (or folding) is valid only between active learning times with at most one of the learning times included. If the process of learning is not constrained to a bounded time interval, the unfolding cannot be globally invariant in time. Furthermore, this proof underlines that input-output functions with different input-domain cannot be functional equivalent.*

**Corollary 3.5** (No Finite Family of FNNs Can Track a Plastic RNN)
*Let* $\mathcal{R}$ *be an RNN with input space* $\mathcal{X}_{\mathcal{R}}$ *and* $\phi_t^{\mathcal{R}} : (\mathcal{X}_{\mathcal{R}})^t \to \mathcal{Y}$ *the input-output map at time* $t$*. Suppose that for every* $T \in \mathbb{N}$ *with* $T > 0$*, there exist* $t_1, t_2 \in \mathbb{N}$ *with* $t_1, t_2 > T$ *and* $t_1 < t_2$ *such that* $(\mathcal{X}_{\mathcal{R}})^{t_1} \subsetneq (\mathcal{X}_{\mathcal{R}})^{t_2}$*. Then, for any finite collection of FNNs* $\{\mathcal{F}_1, \ldots, \mathcal{F}_N\}$*, there does not exist a time* $t^* \geq 0$ *such that, for all* $t \geq t^*$*, there exists* $i \in \{1, \ldots, N\}$ *with* $\mathcal{R} \equiv_t \mathcal{F}_i$*.*

*Proof.* Suppose to the contrary that there exists $t^* \geq 0$ and a finite collection of static FNNs $\{\mathcal{F}_1, \ldots, \mathcal{F}_N\}$ such that for all $t \geq t^*$, there is $i \in \{1, \ldots, N\}$ with $\mathcal{R} \equiv_t \mathcal{F}_i$. This means that, for each $t \geq t^*$, the input domain of $\mathcal{F}_i$ must match that of $\mathcal{R}$ at time $t$: $\mathcal{X}_{\mathcal{F}_i} = (\mathcal{X}_{\mathcal{R}})^t$. But each $\mathcal{F}_i$ has a fixed (time-independent) input domain, and by assumption, for any $T$, there exist $t_1, t_2 > T$, $t_1 < t_2$, such that $(\mathcal{X}_{\mathcal{R}})^{t_1} \subsetneq (\mathcal{X}_{\mathcal{R}})^{t_2}$, so $(\mathcal{X}_{\mathcal{R}})^{t_1} \neq (\mathcal{X}_{\mathcal{R}})^{t_2}$. Since there are only finitely many FNNs, their (fixed) input domains correspond to at most $N$ distinct sets, but for all $t \geq t^*$, there must be an $i$ such that $\mathcal{X}_{\mathcal{F}_i} = (\mathcal{X}_{\mathcal{R}})^t$. Since there are infinitely many $t$ and their domains are distinct for different $t$, this is impossible by the pigeonhole principle. Therefore, it is impossible to choose a finite family of static FNNs so that for all $t \geq t^*$, $\mathcal{R} \equiv_t \mathcal{F}_i$ for some $i$. This contradicts our original assumption, and proves the corollary. ∎

**Remark 3.6**
*Corollary 3.5 implies that no finite family of FNNs* $\{\mathcal{F}_1, \ldots, \mathcal{F}_N\}$ *can track all future maps of* $\mathcal{R}$*, no matter how large* $N$ *is taken. The RNN cannot be tracked by switching between finitely many static FNNs for all sufficiently large* $t$*. This result has several important implications:*

1. ***Dimensional mismatch****: A learning RNN with unbounded effective learning traces out a trajectory in function space that cannot be contained in the finite-dimensional span of any finite collection of static functions.*

2. ***Fundamental limitation for model compression****: This shows that plastic RNNs cannot be exactly replaced by switching between a finite library of static networks, placing theoretical limits on certain model distillation and compression techniques.*

3. ***Distinction from approximation**: While finite collections of FNNs may approximate the RNN's behavior over finite time windows, exact functional equivalence cannot be maintained indefinitely through any finite switching strategy.*

*We will expand on these considerations in the following sections.*

**Remark 3.7**
*The impossibility result in Corollary 3.5 is particularly strong because it holds regardless of how the switching between FNNs is orchestrated, even with perfect knowledge of when to switch and to which network, no finite collection suffices.*

**Remark 3.8** (Learning Differences)
*Consider a FNN $\mathcal{F}_1$ that approximates a RNN $\mathcal{R}$ at time $t_1$ and the FNN $\mathcal{F}_2$ that approximates $\mathcal{R}$ at a following time $t_2 > t_1$. In this situation $\mathcal{F}_2$ must have one extra layer than $\mathcal{F}_1$. However, the weights of the first layers of $\mathcal{F}_2$ do not coincide with those of $\mathcal{F}_1$ because they have to match the RNN at different times.*

**Lemma 3.9** (Asymmetry of Folding and Unfolding)
*Let $\mathcal{F}$ be a feedforward neural network of depth $L$, and let $\mathcal{R}$ be a recurrent neural network.*

1. *There exist a recurrent neural network $\mathcal{R}_\mathcal{F}$ and an encoding $\iota$ such that the first $L$ steps of $\mathcal{R}_\mathcal{F}$ simulate $\mathcal{F}$, i.e.*

$$\phi_L^{\mathcal{R}_\mathcal{F}}(\iota(x)) = \phi^{\mathcal{F}}(x) \qquad \forall x \in \mathcal{X}_F.$$

2. *The network $\mathcal{R}_\mathcal{F}$ may be chosen so that, for times $t > L$, its hidden-state evolution depends non-trivially on subsequent inputs.*

3. *For every $T \in \mathbb{N}$, Theorem 2.1 yields an FNN $\mathcal{F}_T$ such that*

$$\phi^{\mathcal{F}_T} = \phi_T^{\mathcal{R}}$$

*on $(\mathcal{X}_R)^T$.*

4. *If $\mathcal{R}$ satisfies the hypotheses of Theorem 3.2, then no single FNN, and by Corollary 3.5 no finite family of FNNs, can track $\mathcal{R}$ for all sufficiently large times.*

*Thus folding embeds a bounded feedforward computation into an ongoing recurrent process, whereas unfolding yields only horizon-indexed feedforward surrogates. Under the hypotheses of Theorem 3.2, this asymmetry is strict.*

*Proof.* (1) Let $\mathcal{F}$ have depth $L$. Define $\mathcal{R}_\mathcal{F}$ so that its hidden state stores a layer counter and the current layer activation, and let $\iota(x)$ be the length-$L$ encoded input sequence that presents $x$ to this simulation. By construction, the first $L$ recurrent updates of $\mathcal{R}_\mathcal{F}$ simulate the $L$ layers of $\mathcal{F}$, hence

$$\phi_L^{\mathcal{R}_\mathcal{F}}(\iota(x)) = \phi^{\mathcal{F}}(x) \qquad \forall x \in \mathcal{X}_F.$$

(2) The update rule of $\mathcal{R}_\mathcal{F}$ may be defined so that after the first $L$ steps, where it simulates $\mathcal{F}$, it follows any prescribed nontrivial recurrent continuation. This leaves the identity in (1)

unchanged, but yields a recurrent process that can continue beyond depth $L$ and, if desired, depend nontrivially on later inputs.

(3) This is exactly Theorem 2.1: for each fixed $T$ there exists an FNN $\mathcal{F}_T$ such that

$$\phi^{\mathcal{F}_T} = \phi_T^{\mathcal{R}}$$

on $(\mathcal{X}_R)^T$.

(4) This is exactly Theorem 3.2, together with Corollary 3.5: under those hypotheses, no single FNN, and indeed no finite family of FNNs, can track $\mathcal{R}$ for all sufficiently large times.

Therefore folding embeds a bounded feedforward computation into an ongoing recurrent process, whereas unfolding yields only horizon-indexed feedforward surrogates. Under the hypotheses of Theorem 3.2, the asymmetry is strict. ■

### 4 Argument: Learning Implies More Information

The distinction between static FNNs and learning RNNs is fundamental in understanding the expressive power and information content of neural models on sequential data. A static FNN, with fixed or randomly-initialized weights, processes a finite window of inputs via a memoryless mapping, unable to adapt or integrate information across longer temporal contexts. A plastic RNN learns to aggregate information over arbitrarily long sequences through the evolution of its hidden state.

From an information-theoretic perspective, this means that the mutual information between a target variable $Y$ and the state of a learning RNN, $I(Y; h_t)$, can be strictly greater than the mutual information $I(Y; f(X_{t-m+1}, ..., X_t))$ extractable by an FNN on a window of size $m$. The monotonicity of mutual information, together with the Markov property of the RNN hidden states conditional with respect to the input sequence, underpins this argument: the learned recurrent structure is capable of encoding and compressing a complete, task-relevant summary of the entire history, rather than a truncated snapshot.

This distinction becomes especially important when considering the unfolding argument. In fact, unless these weights are adapted to the temporal structure of the data (i.e., the network undergoes learning), the unfolded FNN cannot, in general, accumulate or utilize long-term dependencies in a meaningful way. It is the process of learning that enables the RNN to transcend the limitations of both fixed FNNs and untrained unfolded architectures, yielding greater information content and improved performance on sequential tasks.

**Definition 4.1** (Mutual Information)
*Let $(X, Y)$ be a pair of random variables with joint probability distribution $p_{X,Y}(x, y)$ and marginal distributions $p_X(x)$ and $p_Y(y)$. The mutual information between $X$ and $Y$ is defined as*

$$I(X; Y) = \iint p_{X,Y}(x, y) \log \left( \frac{p_{X,Y}(x, y)}{p_X(x)p_Y(y)} \right) dx\, dy$$

*when $X$ and $Y$ are continuous, or*

$$I(X; Y) = \sum_x \sum_y p_{X,Y}(x, y) \log \left( \frac{p_{X,Y}(x, y)}{p_X(x)p_Y(y)} \right)$$

*when $X$ and $Y$ are discrete.*

**Lemma 4.2** (Data Processing Inequality)
*Suppose $X \to Y \to Z$ forms a Markov chain (i.e., $X$ and $Z$ are conditionally independent given $Y$). Then the data processing inequality states that:*

$$I(X; Y) \geq I(X; Z),$$

*where $I(\cdot; \cdot)$ denotes mutual information.*

**Theorem 4.3** (Conditional Markov Property of RNN Hidden States)
*Let $\{h_t\}_{t=0}^T$ be the sequence of hidden states in a RNN defined recursively by*

$$h_t = F(h_{t-1}, x_t).$$

*Assume $h_0$ is given. Then, for any $t \in \{1, \ldots, T\}$, the sequence $\{h_t\}$ satisfies the conditional (to $\{x_t\}_t$) Markov property: for any measurable set $A \subseteq \mathcal{H}$,*

$$P(h_t \in A \mid h_{t-1}, h_{t-2}, \ldots, h_0, x_1, \ldots, x_T) = P(h_t \in A \mid h_{t-1}, x_t).$$

*Proof.* We have

$$P(h_t \in A \mid h_{t-1}, h_{t-2}, \ldots, h_0, x_1, \ldots, x_T) = P(F(h_{t-1}, x_t) \in A \mid h_{t-1}, h_{t-2}, \ldots, h_0, x_1, \ldots, x_T)$$

$$= P(F(h_{t-1}, x_t) \in A \mid h_{t-1}, x_t) = P(h_t \in A \mid h_{t-1}, x_t).$$

This is precisely the definition of the (conditional to $\{x_t\}_t$) Markov property. ■

**Remark 4.4**
*This theorem works also in the case in which the sequence $\{x_t\}_{t \in \mathbb{N}}$ is deterministic.*

**Theorem 4.5** (Monotonicity of Mutual Information Under Measurable Processing)
*Let $(\Omega, \mathcal{F}, P)$ be a probability space. Let $Y : \Omega \to \mathcal{Y}$ be a random variable, and $T : \Omega \to \mathcal{T}$, $S : \Omega \to \mathcal{S}$ be random vectors defined on $\Omega$ with $S = \pi(T)$ for some measurable projection $\pi : \mathcal{T} \to \mathcal{S}$. Let $g : \mathcal{T} \to \mathcal{Z}_g$ and $f : \mathcal{S} \to \mathcal{Z}_f$ be measurable functions such that $f = h \circ g \circ \iota$ for some measurable $h : \mathcal{Z}_g \to \mathcal{Z}_f$ and measurable inclusion $\iota : \mathcal{S} \hookrightarrow \mathcal{T}$ (i.e., $f(S) = h(g(T))$ whenever $S = \pi(T)$). Then,*

$$I(Y; g(T)) \geq I(Y; f(S)),$$

*where $I(\cdot;\cdot)$ denotes the mutual information under $P$.*

*Proof.* By the data processing inequality ( Theorem 2.8.1 in [1]), for any measurable function $h$,

$$I(Y; g(T)) \geq I(Y; h(g(T))) = I(Y; f(S)),$$

since $f(S) = h(g(T))$ almost surely by construction. ■

**Remark 4.6** (Counterexample (Necessity of $f = h \circ g$))
*Let $T = (X_1, X_2)$ and $S = X_1$. Define*

$$f(S) = f(X_1) = X_1, \qquad g(T) = g(X_1, X_2) = X_2.$$

*Let $Y = X_1$, where $X_1$ and $X_2$ are independent random bits, each uniformly distributed on $\{0, 1\}$. Then*

$$I(Y; f(S)) = I(X_1; X_1) = H(X_1) = 1,$$

*while*

$$I(Y; g(T)) = I(X_1; X_2) = 0,$$

*since $X_1$ and $X_2$ are independent. Note that $f$ cannot be written as $h \circ g$ for any function $h$ (i.e., $X_1$ is not a function of $X_2$). In this case,*

$$I(Y; f(S)) > I(Y; g(T)).$$

*Thus, the monotonicity $I(Y; g(T)) \geq I(Y; f(S))$ does not hold if $f$ is not a function of $g$.*

**Corollary 4.7**
*In the framework of Theorem 4.5, let $S = (X_{t-m+1}, \dots, X_t)$ be a length-$m$ window of a time series, $T$ a set containing $S$ (e.g., $T = (X_1, X_2, \dots, X_t)$), $\phi^{\mathcal{F}}$ being the input-output function of a static FNN with input $S$, and $\phi_t^{\mathcal{R}}$ the output of a RNN, with the first $m$-layers ($m < t$) coinciding with the FNN. Then*

$$I(Y; g(T)) \geq I(Y; f(S)),$$

*where $Y$ is any target variable.*

*Proof.* By assumption, $S \subseteq T$, and the RNN $g(T)$ contains all the information that $f(S)$ uses, since its first $m$ layers coincide with $f$ on $S$. There exists a measurable function $h$ such that $f(S) = h(g(T))$: simply restrict $g(T)$ to its output after processing the first $m$ components (since these coincide with $f$), and ignore any subsequent computation. Thus, $f = h \circ g$. By Theorem 4.5 and Lemma 4.2,

$$I(Y; f(S)) \leq I(Y; g(T)).$$

■

**Remark 4.8**
*Note that this proof does not cover the case of a fixed FNN with randomly initiated weights, which can, by chance, have more information about the output $Y$ that an evolving RNN at time $t$. This happens with probability zero. This statement can be made rigorous, but it would be very technical and the development of the needed complicated machinery goes beyond the scope of this paper.*

**Remark 4.9** (Information Value of Learning)
*A static FNN defines a map from a fixed window of $m$ consecutive inputs $S = (X_{t-m+1}, \dots, X_t)$ to an output through $\phi^{\mathcal{F}}$, with no explicit memory or mechanism for aggregation of dependencies beyond this window. In contrast, a learning RNN generates a map $\phi_t^{\mathcal{R}}$ from the full sequence $T = (X_1, \dots, X_t)$ to an output, allowing for long-term dependence. Theorem 4.3 established above guarantees that, conditioned on the entire input sequence, the hidden states summarize all relevant information from the entire past up to time $t$. By Theorem 4.5, a measurable function (such as a sufficiently expressive RNN) operating on a superset of the data (such as all past inputs) can, in principle, encode at least as much, and typically more, information about a target variable $Y$ than an FNN with access solely to the window $S$.*

### 5 Argument: Learning at Discrete Times Needs Countable Unfolding

It is a foundational insight of the unfolding theorem [2] that a RNN, when unrolled for a finite number $T$ of discrete time steps, can be represented as a FNN of depth $T$ with parameter sharing across layers (See Argument: Learning Limits Unfolding). This correspondence, however, crucially hinges on the fact that time is discretized and the sequence length is finite. When learning proceeds at discrete times, the process naturally induces a *countable* unfolding: each time step corresponds to a new layer in the unrolled FNN, and thus the total depth of the equivalent FNN grows linearly with the number of discrete learning iterations. As a result, for an RNN trained or evaluated over arbitrarily long (but countable) temporal horizons, the theoretically equivalent FNN would require a countable, potentially infinite, number of layers to capture the same iterative computation. Therefore, the act of learning at discrete times is inherently linked to countable unfolding, illuminating both the expressive power and the limitations that arise from approximating recurrent computations with static, unrolled architectures. We want to give a discrete dynamical systems perspective of this.

**Definition 5.1** (Discrete Dynamical System)
*A (discrete-time) dynamical system is a tuple*

$$\mathcal{D} = (\mathcal{S}, F),$$

*where $\mathcal{S}$ is the state space and $F : \mathcal{S} \to \mathcal{S}$ is the state transition map. Given $s_0 \in \mathcal{S}$, the system evolves as*

$$s_{t+1} = F(s_t).$$

*The sequence $(s_t)_{t\geq 0}$ is called the* orbit *starting at $s_0$. A state $s^* \in \mathcal{S}$ is a* fixed point *if $F(s^*) = s^*$.*

**Theorem 5.2** (Fixed Point - No Learning)
*Let $\mathcal{D} = (\mathcal{S}, F)$ be a discrete dynamical system. If $F : \mathcal{S} \to \mathcal{S}$ is defined by $F(s) = s$ for all $s \in \mathcal{S}$, then every $s^* \in \mathcal{S}$ is a fixed point of $\mathcal{D}$.*

*Proof.* By definition, a state $s^* \in \mathcal{S}$ is called a fixed point of $\mathcal{D}$ if $F(s^*) = s^*$. For a static FNN, by assumption,

$$F(s) = s \quad \text{for all} \quad s \in \mathcal{S}.$$

In particular, for any $s^* \in \mathcal{S}$,

$$F(s^*) = s^*.$$

Therefore, $s^*$ is a fixed point of $F$. Since $s^*$ was arbitrary, it follows that every state in $\mathcal{S}$ is a fixed point. ∎

**Remark 5.3**
*The case $F = Id$ represents the case of no learning, as the state is never updated.*

**Corollary 5.4** (Static FNN as a Fixed Point - No Learning)
*Consider the dynamical system $\mathcal{D} = (\mathcal{S}, F)$, with $\mathcal{S} = C(\mathcal{X}, \mathcal{Y})$ the space of continuous functions with $(\mathcal{X}, d_{\mathcal{X}})$ a compact metric space and $(\mathcal{Y}, \|\cdot\|_{\mathcal{Y}}$ a normed space. Suppose $F = I$, the identity map in $C(\mathcal{X}, \mathcal{Y})$. Then, if $s_0 = \phi^{\mathcal{F}}$ is the input-output function of a FNN $\mathcal{F}$, then $F^k(s_0) = s_0$ for every $k \in \mathbb{N}$ with $F^k := F \circ \cdots \circ F$, composed $k$ times.*

*Proof.* This is a direct application of Theorem 5.2, with $\mathcal{S} = C(\mathcal{X}, \mathcal{Y})$ and $F = \mathrm{Id}_{C(\mathcal{X},\mathcal{Y})}$. By the theorem, since $F(f) = f$ for all $f \in C(\mathcal{X}, \mathcal{Y})$. In particular, if $\phi^{\mathcal{F}} \in C(\mathcal{X}, \mathcal{Y})$ is the input-output map of static FNN, then $F(\phi^{\mathcal{F}}) = \phi^{\mathcal{F}}$. ■

**Remark 5.5**
*The absence of learning ($F = \mathrm{Id}_{C(\mathcal{X},\mathcal{Y})}$) implies that any network, and so also an FNN, is a fixed point of the dynamical system.*

**Proposition 5.6** (Plastic RNN as an Orbit - Unbounded Learning)
*Consider the dynamical system $\mathcal{D} = (\mathcal{S}, F)$, with $\mathcal{S} = C(\mathcal{X}, \mathcal{Y})$ the space of continuous functions with $(\mathcal{X}, d_{\mathcal{X}})$ a compact metric space and $(\mathcal{Y}, \|\cdot\|_{\mathcal{Y}}$ a normed space. Suppose $F^l(s^*)) \neq s^*$ for every $s^*$ in $C(\mathcal{X}, \mathcal{Y})$ and $l \in \mathbb{N}$. Let $s_0 = \phi^{\mathcal{R}_0}$ be the input-output function of a RNN $\mathcal{R}$. Then, $\{F^k(s_0)\}_{k\in\mathbb{N}}$ is a countable set with $F^k := F \circ \cdots \circ F$, composed $k$ times.*

*Proof.* By hypothesis, $F^k(s_0) \neq s_0$ for every $k \in \mathbb{N}$ with $F^k := F \circ \cdots \circ F$, composed $k$ times. Therefore, $\{F^k(s_0)\}_{k\in\mathbb{N}}$ is a countable set. ■

**Remark 5.7**
*The presence of perpetual learning ($F^l(s^*)) \neq s^*$ for every $s^*$ in $C(\mathcal{X}, \mathcal{Y})$ and $l \in \mathbb{N}$) implies that an RNN draws a discrete orbit of the dynamical system.*

**Corollary 5.8** (Contrast Between FNN and RNN)
*A feedforward network is equivalent to evaluating a finite recursion that halts in a fixed output; hence it corresponds to a* fixed point *of a computation. A recurrent network is equivalent to a recurrent state-update rule; hence it generates an* orbit *of hidden states in a dynamical system.*

**Proposition 5.9**
*Let $\mathcal{F}$ be a FNN representing the target function $f^* = \phi^{\mathcal{F}_L}$, and let $\mathcal{R}$ be a RNN with learning dynamics generating a trajectory (orbit) of input-output maps*

$$\{\phi_t^{\mathcal{R}}\}_{t\geq 0}.$$

*Suppose the RNN is trained to approximate $f^*$. Then:*

1. *If for all $t$, $\phi_t^{\mathcal{R}} \neq f^*$ (i.e., at no time does the RNN exactly replicate the FNN), then $\mathcal{F}$ is a better approximator of $f^*$ than $\mathcal{R}$ at every time $t$.*
2. *If there exists $t^*$ such that $\phi_{t^*}^{\mathcal{R}} = f^*$, then at that time the approximating power of $\mathcal{R}$ matches $\mathcal{F}$.*

*In particular, the static FNN is a better approximator of itself than any learning RNN whose orbit never passes through $f^*$.*

*Proof.* Let $f^*$ be the function realized by the static FNN. Let $\left\{\phi_t^{\mathcal{R}} : \mathcal{X}_R^t \to \mathcal{Y}_R\right\}_{t\geq 0}$ be the sequence of input-output maps (the orbit) produced by the learning RNN as it updates its parameters. Assume the goal is to approximate $f^*$.

**Case 1:** Suppose for all $t$, $\phi_t^{\mathcal{R}} \neq f^*$. Then, at all times, the RNN does not exactly realize $f^*$. Since $\mathcal{F}$ by definition always realizes $f^*$, for any input $x$,

$$\phi^{\mathcal{F}_L}(x) = f^*(x),$$

while for $\mathcal{R}$,

$$\forall t : \phi_t^{\mathcal{R}}(x) \neq f^*(x).$$

Thus, pointwise (and hence uniformly),

$$\sup_x |\phi^{\mathcal{F}_L}(x) - f^*(x)| = 0 < \sup_x |\phi_t^{\mathcal{R}}(x) - f^*(x)|,$$

so the FNN is a strictly better approximator of itself.

**Case 2:** If there exists $t^*$ such that $\phi_{t^*}^{\mathcal{R}} = f^*$, then at time $t^*$, both the FNN and the RNN realize $f^*$, and thus their approximation error matches (both zero).

■

**Remark 5.10**
*This theorem states that a static FNN is better than a RNN to approximate itself in the case the learning RNN does not encounter the FNN in one of the iterates.*

### 6 Argument: Learning Implies Perturbation Response

A central challenge in the theory of consciousness is to explain how a system can maintain continuous conscious experience that integrates information across time, responding adaptively to both immediate and past events. This challenge is especially salient when contrasting static, feedforward models, often used in traditional computational theories of mind and in static formulations like Integrated Information Theory (IIT), with dynamic, recurrent, or learning-based models that possess internal memory and temporally extended state [3]. The *unfolding argument* formalizes the idea that the dynamical unfolding of recurrent computations cannot, in general, be faithfully simulated by any finite static feedforward network without loss of key properties, such as the long-term persistence of information or the integration of temporally separated events.

Within this theoretical landscape, learning systems such as RNNs exhibit a fundamental sensitivity to localized perturbations: a single, momentary alteration in input can have enduring influence on the hidden state and subsequent output, reflecting an inherent temporal memory and adaptivity. In contrast, static FNNs process input in fixed windows, and any effect of past perturbations is rapidly "forgotten" once outside this window. This distinction has crucial implications for consciousness. If conscious experience requires the ability to integrate and respond to information from the distant past, persistently carrying forward the causal footprint of events, then only systems that support such temporal persistence (as in learning RNNs) can satisfy this requirement. Static, window-limited models fall short, as their lack of persistent effect beyond a finite input window structurally excludes true temporal integration. Thus, the ability of a system to exhibit different responses to localized perturbations, and for those differences to persist under unfolding, provides a theoretical and experimentally accessible signature distinguishing genuinely dynamical from static architectures, and by extension, marks a core difference between dynamical and static theories of consciousness.

The following argument makes this distinction precise, laying a rigorous mathematical foundation for why learning systems, by virtue of their temporal dynamism, are uniquely equipped to model the persistent and integrated nature of conscious experience, while static systems inherently forget.

**Theorem 6.1** (Insensitivity of Static FNNs to Localized Perturbations Outside Input Window)
*Let $(x_t)_{t\in\mathbb{N}}$ be a time series with values in $\mathcal{X} \subset \mathbb{R}^n$. Fix $m \in \mathbb{N}$ and let*

$$\phi^{\mathcal{F}_L} : \mathcal{X}_{F_L} \to \mathcal{Y}_{F_L}$$

*be a (measurable) input-output function of a FNN $\mathcal{F}$ mapping $m$-length input windows to outputs. For each $t \geq m-1$, define the network output*

$$y_t = F(x_{t-m+1}, x_{t-m+2}, \ldots, x_t).$$

*Now, consider a perturbed time series $\tilde{x}_s = x_s$ for all $s \neq k$, and $\tilde{x}_k = x_k + \delta$ for some $k \geq 0$ and $\delta \in \mathcal{X}$. Then for any $t \geq k+m$, the outputs satisfy*

$$y_t = \tilde{y}_t,$$

*where $\tilde{y}_t$ is the output for the perturbed series. In other words, after $m$ steps, the effect of any perturbation at time $k$ is forgotten, and the output is independent of $\delta$:*

$$\phi^{\mathcal{F}_L}(\tilde{x}_{t-m+1},\ldots,\tilde{x}_t) = \phi^{\mathcal{F}_L}(x_{t-m+1},\ldots,x_t), \quad \forall t \geq k+m.$$

*Proof.* Let $\mathbf{x} = (x_t)_{t\in\mathbb{N}}$ be a time series, and let $\tilde{\mathbf{x}}$ be the perturbed series with $\tilde{x}_s = x_s$ for $s \neq k$ and $\tilde{x}_k = x_k + \delta$. By the definition of the sliding-window static FNN, for any $t \geq m-1$,

$$y_t = \phi^{\mathcal{F}_L}(x_{t-m+1}, x_{t-m+2},\ldots,x_t), \qquad \tilde{y}_t = \phi^{\mathcal{F}_L}(\tilde{x}_{t-m+1},\ldots,\tilde{x}_t).$$

For $t \geq k+m$, observe that the window $(t-m+1)$ to $t$ lies entirely after $k$:

$$t-m+1 > k \implies t > k+m-1 \implies t \geq k+m,$$

since $t$ and $k$ are integers. Therefore, for all $s \in [t-m+1, t]$ when $t \geq k+m$, we have $s > k$, so $\tilde{x}_s = x_s$ (the perturbation $\delta$ affects only $\tilde{x}_k$). Thus,

$$(\tilde{x}_{t-m+1},\ldots,\tilde{x}_t) = (x_{t-m+1},\ldots,x_t)$$

and consequently,

$$\tilde{y}_t = F(\tilde{x}_{t-m+1},\ldots,\tilde{x}_t) = F(x_{t-m+1},\ldots,x_t) = y_t,$$

for all $t \geq k+m$. Therefore, the effect of the perturbation at time $k$ vanishes after $m$ steps, as claimed. ∎

**Theorem 6.2** (Persistence of Pertubation Effect in RNNs)
*Let $(x_t)_{t\in\mathbb{N}}$ be an input sequence with $x_t \in \mathcal{X} \subseteq \mathbb{R}^n$. Consider a RNN defined by an initial state $h_0 \in \mathcal{H}$ and update rules*

$$h_t = f(h_{t-1}, x_t), \qquad y_t = g(h_t),$$

*where $f : \mathcal{H}_R \times \mathcal{X}_R \times \mathcal{W}_R \to \mathcal{H}_R$ ang $g : \mathcal{H}_R \times \mathcal{X}_R \times \mathcal{W}_R \to \mathcal{Y}_R$ are measurable functions. Fix $k \in \mathbb{N}_{\geq 1}$ and a perturbation $\delta \in \mathcal{X}_R \setminus \{0\}$. Define the perturbed input sequence $(\tilde{x}_t)$ by*

$$\tilde{x}_s = \begin{cases} x_k + \delta, & s = k, \\ x_s, & s \neq k. \end{cases}$$

*Let $(\tilde{h}_t, \tilde{y}_t)$ be the hidden states and outputs of the RNN driven by $(\tilde{x}_t)$, starting from the same $h_0$. Then for each $t \geq k$,*

$$h_t \neq \tilde{h}_t \quad \implies \quad y_t \neq \tilde{y}_t,$$

*unless one of the following degeneracies occurs:*

1. ***Invariance in hidden dynamics:***

   $$f(h_{k-1}, x_k) = f(h_{k-1}, x_k + \delta),$$

   *or, more generally, $f(h, x) = f(h, x+\delta)$ for the relevant arguments; or*

*2.* ***Invariance in output mapping:*** *$g(h_t) = g(\tilde{h}_t)$ despite $h_t \neq \tilde{h}_t$.*

*In particular, in the absence of such degeneracies (e.g., in generic settings where $f$ and $g$ are non-degenerate smooth maps), the effect of the perturbation $\delta$ at time $k$ induces a difference*

$$y_t - \tilde{y}_t \neq 0, \qquad \forall t \geq k,$$

*so that the perturbation is not forgotten in finite time.*

*Proof.* Let $(x_t)_{t\in\mathbb{N}}, (h_t)_{t\in\mathbb{N}}, (y_t)_{t\in\mathbb{N}}$ be the original input, hidden state, and output sequences, defined by

$$h_0 \in \mathcal{H}_R, \qquad h_t = f(h_{t-1}, x_t), \qquad y_t = g(h_t).$$

Let $(\tilde{x}_t)_{t\in\mathbb{N}}, (\tilde{h}_t)_{t\in\mathbb{N}}, (\tilde{y}_t)_{t\in\mathbb{N}}$ be the perturbed input, hidden state, and output sequences, where for some fixed $k \in \mathbb{N}_{\geq 1}$ and $\delta \in \mathcal{X}_R \setminus \{0\}$,

$$\tilde{x}_s = \begin{cases} x_k + \delta, & s = k, \\ x_s, & s \neq k, \end{cases}$$

with $\tilde{h}_0 = h_0$, and for $t \geq 1$:

$$\tilde{h}_t = f(\tilde{h}_{t-1}, \tilde{x}_t), \qquad \tilde{y}_t = g(\tilde{h}_t).$$

We prove by induction on $t \geq k$ that unless at least one degenerate condition of the theorem holds, a single perturbation at time $k$ induces a difference in $(h_t)$ and $(\tilde{h}_t)$, and thus in $(y_t)$ and $(\tilde{y}_t)$, for all future $t$.

**Base case ($t = k$):**

$$\begin{aligned} h_k &= f(h_{k-1}, x_k), \\ \tilde{h}_k &= f(h_{k-1}, x_k + \delta). \end{aligned}$$

If $f(h_{k-1}, x_k) \neq f(h_{k-1}, x_k + \delta)$ (i.e., $f$ is not invariant to the perturbation at $(h_{k-1}, x_k)$ and $\delta \neq 0$), then $h_k \neq \tilde{h}_k$. If, in addition, $g(h_k) \neq g(\tilde{h}_k)$ (i.e., $g$ is not invariant to differences in its argument), then $y_k \neq \tilde{y}_k$.

**Inductive step:** Suppose for some $t \geq k$ we have $h_t \neq \tilde{h}_t$. Consider $t + 1$:

$$h_{t+1} = f(h_t, x_{t+1}), \qquad \tilde{h}_{t+1} = f(\tilde{h}_t, x_{t+1}).$$

If $f(h, x)$ is sensitive to changes in its first argument, $h_{t+1} \neq \tilde{h}_{t+1}$. Otherwise, if $f$ is invariant to changes in $h$ for the input $x_{t+1}$ and the current $h_t$, then degeneration (1) from the theorem statement has occurred. By identical reasoning, if $g$ is injective (not output-invariant for $h_{t+1} \neq \tilde{h}_{t+1}$), then $y_{t+1} \neq \tilde{y}_{t+1}$. Otherwise, degeneration (2) from the theorem statement occurs. Thus, provided neither degenerate case holds at any time $t \geq k$, the effect of the input perturbation $\delta$ at time $k$ persists indefinitely in the hidden states and, unless masked by $g$, in the outputs. Formally,

$$y_t - \tilde{y}_t \neq 0 \quad \forall\, t \geq k,$$

unless there exists $t \geq k$ such that either

$$f(h_{t-1}, x_t) = f(\tilde{h}_{t-1}, x_t) \quad \text{or} \quad g(h_t) = g(\tilde{h}_t).$$

This concludes the proof. ■

**Corollary 6.3** (Persistence of Perturbation Effect under Unfolding in RNNs)
*Let* $(x_t)$ *and* $k, \delta$ *be as above, and consider the* $T$*-step unfolded version of a trained RNN, i.e., a composition of* $T$ *instances of* $(f, g)$*. Then, for any* $t \geq k$ *with* $t - k < T$*, the conclusions of Theorem 6.2 hold: the output difference* $y_t - \tilde{y}_t$ *remains (in general) nonzero unless degeneracy occurs.*

*Proof.* Follows from Theorem 6.2 and Definitions 1.1 and 1.2. ■

**Remark 6.4**
*Experimental Consequences. Theorem 6.2 and Corollary 6.3 demonstrate that it is possible to experimentally validate the persistence of localized perturbations in RNNs by comparing outputs on original and perturbed input sequences, even when using an unfolded computational graph, as long as the number of unrolled steps* $T$ *is at least as large as the time horizon of interest after the perturbation at* $k$*. By contrast, Theorem 6.1 shows that for static FNNs, the effect of a single perturbation is strictly confined by the input window size and is forgotten after* $m$ *steps. There is no such forgetting in the generic RNN setting (although vanishing gradient might occur [2]). Therefore, experimental protocols that probe the effect of input perturbations in learning RNNs remain sensitive to the persistence phenomenon. The theoretical distinction between static FNNs and RNNs is preserved under unfolding, and the persistence of perturbation effects can be tested in actual experiments on trained RNNs, by comparing outputs on original and perturbed inputs.*

*Our theorems demonstrate that experimental protocols comparing original and perturbed input sequences in RNNs, even when implemented by unfolding, are valid for probing persistence effects, crucial for memory, temporal integration, and thus for any consciousness theory aspiring to explain continuity of awareness. Static computational models, including standard FNNs and static formulations of IIT, do not support these phenomena. As such, any static theory of consciousness that does not explicitly account for persistent, temporally unified states may fail to capture essential features of conscious experience.*

### 7 Argument: Learning Implies Approximation Capacity

*In Section 3, we showed that learning limits the unfolding/folding argument, in the sense that unfolding/folding is possible to the same network only in a time interval with one single learning episode at the beginning of the time interval. In Section 3, we do not address the problem on how learning is the key to approximation capacity. Without learning, namely with a static network, there is essentially no approximation capacity, as the neurons are not allowed to train and adapt to the task at hand. Approximation is a metric concept and so, with respect to Section 3, we need to introduce extra machinery in order to have a way to measure distance between functions. Recall that, in infinite dimensional spaces, such as any functional space without low-dimensional constraints (e.g. smooth functions vs polynomials or order less than 3), norms are not equivalent and so any approximation theory in function spaces is sensitive to the norm used [4]. We will not enter in these subtleties here and we will concentrate on the continuous norm, as typically used in classical statistical learning theory arguments [5]. Despite this, the fact that learning is necessary for unbounded approximation capacity, is not affected. Several results on universal approximation properties of RNNs are available in the literature. We refer to [6] for the precise statement of the theorem and previous results in the field.*

**Definition 7.1** ($L^\infty$ Distance Between Functions)
*Let $\mathcal{F} \subseteq C(\mathcal{X}, \mathcal{Y})$ be a family of functions from a compact metric space $(\mathcal{X}, d_\mathcal{X})$ to a normed space $\mathcal{Y}$. For $f, g \in \mathcal{F}$, define*

$$\|f - g\|_{L^\infty(\mathcal{X})} := \sup_{x \in \mathcal{X}} \|f(x) - g(x)\|_\mathcal{Y}.$$

**Theorem 7.2** (Static Networks Can Only Approximate Themselves)
*Let $(\mathcal{X}, d_\mathcal{X})$ be a compact metric space and $(\mathcal{Y}, \|\cdot\|_\mathcal{Y})$ a normed space. Let $f : \mathcal{X} \to \mathcal{Y}$ be the input-output function realized by a fixed (FNN) of depth $L$:*

$$\mathcal{F} = (\{\mathcal{H}_\ell\}_{\ell=1}^L, \{W_{F,\ell}\}_{\ell=1}^L, \{f_\ell\}_{\ell=1}^L)$$

.
*Then for any continuous target function $g \in C(\mathcal{X}, \mathcal{Y})$, either*

$$g = f,$$

*or there exists $\epsilon_0 > 0$, such that for $\epsilon < \epsilon_0$*

$$\|f - g\|_{L^\infty(\mathcal{X})} > \epsilon.$$

*Proof.* Since $\mathcal{X}$ is compact and $f, g$ are continuous, the difference

$$d(f, g) := \sup_{x \in \mathcal{X}} \|f(x) - g(x)\|_\mathcal{Y}$$

is finite and attained. If $g = f$, then $d(f, g) = 0$. If $g \neq f$, then there exists some $x_0 \in \mathcal{X}$ such that $f(x_0) \neq g(x_0)$. By continuity,

$$\|f(x_0) - g(x_0)\|_\mathcal{Y} > 0.$$

Hence $d(f, g) \geq \|f(x_0) - g(x_0)\|_{\mathcal{Y}} > 0$. In particular, taking $\epsilon_0 := d(f, g)/2$, we see that

$$\|f - g\|_{L^\infty(\mathcal{X})} = \epsilon > 0,$$

for every $\epsilon < \epsilon_0$. Thus, the fixed FNN can approximate no function except itself. ∎

**Remark 7.3**
*Theorem 7.2 implies that the FNN can only approximate itself and it does it exactly. Otherwise, it will always stay at a positive minimal distance from the function to be approximated.*

**Remark 7.4** (Learning Enables Approximation of Arbitrary Functions)
*Unlike the static FNN that can only realize a single function, the learning RNN can approximate arbitrary target functions on compact domains. In contrast to ongoing learning, during which the approximation capacity can be made arbitrarily large [6], any finite or countable union of static (without learning) FNNs yields a function family with bounded approximation capacity. Note that this is okay, only in the case you want to approximate the functions that are part of this family or generated by them. More generally, for a finite family of FNNs (or, more generally, any class of functions with finite cardinality $|\mathcal{G}| = N < \infty$), there are only a finite number of functions that can be approximated and are those of the family itself, that are matched exactly. In other words, two disjoint finite families of continuous functions on a compact domain are positively separated in $L^\infty$.*

**Theorem 7.5** (Positive $L^\infty$-Separation Between Disjoint Finite Families)
*Let $(\mathcal{X}, d_{\mathcal{X}})$ be a compact metric space and $(\mathcal{Y}, \|\cdot\|_{\mathcal{Y}})$ a normed space. Let $\mathcal{F} = \{f_1, \dots, f_N\} \subseteq C(\mathcal{X}, \mathcal{Y})$ and $\mathcal{G} = \{g_1, \dots, g_M\} \subseteq C(\mathcal{X}, \mathcal{Y})$ be two finite families of continuous functions such that*

$$\mathcal{F} \cap \mathcal{G} = \emptyset.$$

*Then, there exists $\delta > 0$ such that*

$$\|f - g\|_{L^\infty(\mathcal{X})} \geq \delta, \quad \forall f \in \mathcal{F},\ g \in \mathcal{G}.$$

*Proof.* Fix distinct functions $f, g \in C(\mathcal{X}, \mathcal{Y})$. Since $\mathcal{X}$ is compact and $f, g$ are continuous, the difference

$$d(f, g) := \sup_{x \in \mathcal{X}} \|f(x) - g(x)\|_{\mathcal{Y}}$$

is finite and attained at some $x_0 \in \mathcal{X}$. If $f = g$, then $d(f, g) = 0$. If $f \neq g$, there exists $x_0 \in \mathcal{X}$ with $f(x_0) \neq g(x_0)$. Thus $\|f(x_0) - g(x_0)\|_{\mathcal{Y}} > 0$, and by definition

$$d(f, g) = \|f - g\|_{L^\infty(\mathcal{X})} > 0.$$

Therefore, any two distinct functions have strictly positive $L^\infty$-distance. Now consider the finite set of pairwise distances

$$D := \{\|f_i - g_j\|_{L^\infty(\mathcal{X})} : 1 \leq i \leq N,\ 1 \leq j \leq M\}.$$

By assumption, $\mathcal{F} \cap \mathcal{G} = \emptyset$, so $f_i \neq g_j$ for all $(i, j)$. Hence every element of $D$ is strictly positive. Since $D$ is a finite set of positive real numbers, it has a positive minimum:

$$\delta := \min D > 0.$$

By construction,

$$\|f - g\|_{L^\infty(\mathcal{X})} \geq \delta \quad \forall f \in \mathcal{F},\ g \in \mathcal{G}.$$

This proves the claim. ■

**Remark 7.6** (No Adaptation Without Learning: Neurons vs. Environment)
*This theorem can be interpreted with the fact that a static network cannot learn a static environment either, unless the network is already built, at least in part, in that environment, despite the fact that learning is not present. More specifically, let $\mathcal{F} \subseteq C(\mathcal{X}, \mathcal{Y})$ denote the finite family of functions realizable by a static (non-learning) neural system (i.e., the "neurons" or network outputs), and let $\mathcal{G} \subseteq C(\mathcal{X}, \mathcal{Y})$ denote a finite or infinite family of possible environments (tasks, or functions to be realized). Then:*

1. *If there exists $g \in \mathcal{G}$ such that $g \in \mathcal{F}$, the neuronal system can realize this environment exactly (i.e., it is "pre-adapted" to $g$ by built-in design).*

2. *If $g \in \mathcal{G}$ but $g \notin \mathcal{F}$, a static system cannot adapt to or "learn" this environment after deployment. Its response is restricted to the family $\mathcal{F}$ chosen at construction, without the capacity to adjust based on the encountered environment $g$.*

*Therefore, a static collection of neurons cannot adapt to (learn) an environment, unless that environment is already representable among its fixed pre-existing functions.*

| ***Property*** | ***Static Network*** | ***Learning Network*** |
|---|---|---|
| *Adaptation* | *None (fixed)* | *Yes (adaptive)* |
| *Fit to environment* | *Only if predesigned* | *Can adjust/learn* |
| *Limitation* | *No post-design change* | *Limited by update rule* |

**Remark 7.7** (Unfolding Does Not Confer Adaptation to Static Networks)
*The process of unfolding a static (non-learning) network over $T$ steps produces a large feed-forward ("static" unfolded) network whose behavior is entirely determined by its original architecture and fixed parameters. That is, the unfolded computation encodes only those input-output behaviors that were already present at construction; it cannot accommodate, adapt, or learn a new environment encountered after deployment. Genuine adaptation and perpetual learning require dynamic changes of parameters, namely learning rules that update the mapping as new data is encountered. Thus, unfolding, by itself, does not endow a static network with learning or adaptation capabilities.*

**Remark 7.8** (Approximation Capacity and Adaptability)
*Let $\mathcal{F} \subseteq C(\mathcal{X}, \mathcal{Y})$ denote the function class realizable by a neural system (the "neurons"), and let $\mathcal{G} \subseteq C(\mathcal{X}, \mathcal{Y})$ denote the family of possible environments the system may encounter. The* approximation capacity *of $\mathcal{F}$ quantifies how richly the neurons can realize, approximate, or adapt to arbitrary environments in $\mathcal{G}$.*

- *For a* ***static*** *(non-learning) neural system, $\mathcal{F}$ is finite or otherwise fixed, so it can approximate only itself. In this case, only those environments $g \in \mathcal{G}$ which are already contained in $\mathcal{F}$ can be realized with predetermined accuracy. In fact they are learned exactly.*

- *For an **adaptive/learning** system, $\mathcal{F}$ may encompass a much larger, potentially unbounded, space of functions, expanding as the system learns. Here, the neurons can continuously modify their functional output to approximate or match new environments in $\mathcal{G}$, reflecting true capacity for adaptation.*

### 8 Kleiner–Hoel and Lenient Dependency

*Kleiner and Hoel [7] distinguish* inference data *(externally observable input–output behavior) from* prediction data *(the internal organization posited by a theory, such as state variables, connectivity, update rules, memory structure, and plasticity laws). Their core claim is that a theory is empirically testable only if inference data constrains prediction data in a way that is neither vacuous nor uniquely identifying. They call this intermediate regime* lenient dependency*.*

*In the present appendix, the relevant inference data are finite behavioral traces* Beh *obtained under a fixed experimental protocol, possibly including controlled perturbations. The relevant prediction data are the architectural and dynamical features of candidate models, including whether a system is feedforward or recurrent, whether it has bounded memory, and whether it exhibits non-trivial plasticity.*

*Fix a model class $\mathcal{S}$ and an experimental protocol $P$. For each $S \in \mathcal{S}$, let*

$$\mathrm{Beh}_T^P(S) := (x_{1:T}, y_{1:T}^{S,P})$$

*denote the length-$T$ input–output trace produced by $S$ under $P$, where $x_{1:T}$ is the imposed input sequence and $y_{1:T}^{S,P}$ is the resulting output trace. Given observed data*

$$D_T^P := (x_{1:T}, y_{1:T}^{\mathrm{obs}}),$$

*define the* consistency set

$$\mathcal{C}_T^P(D_T^P) := \{\, S \in \mathcal{S} : \mathrm{Beh}_T^P(S) = D_T^P \,\}.$$

*Thus $\mathcal{C}_T^P(D_T^P)$ is the set of candidate internal organizations whose predicted behavior agrees with the observed trace through horizon $T$. For empirical applications, exact equality would typically be replaced by an appropriate tolerance relation or statistical goodness-of-fit criterion; the exact-match formulation is used here only to keep the logical structure transparent.*

**Lemma 8.1** (Consistency sets are nested under horizon extension)
*Let $T < T'$, and suppose $D_{T'}^P$ extends $D_T^P$ by continuing the same experimental protocol. Then*

$$\mathcal{C}_{T'}^P(D_{T'}^P) \subseteq \mathcal{C}_T^P(D_T^P).$$

*Proof.* If a model matches the observed trace through horizon $T'$, then it matches, in particular, the prefix of that trace through horizon $T$. Hence membership in $\mathcal{C}_{T'}^P(D_{T'}^P)$ implies membership in $\mathcal{C}_T^P(D_T^P)$. ■

*The nesting in Lemma 8.1 can be non-trivial:*

**Proposition 8.2** (Extended traces can exclude bounded-memory static surrogates)
*Fix a protocol $P$ and observed data $D_T^P$. Let $S$ be a static feedforward surrogate with finite window size $m$. Suppose that under $P$ the observed behavior exhibits a localized perturbation at some time $k$ whose effect persists at least through some later time $t \geq k + m + 1$. Then*

$$S \notin \mathcal{C}_t^P(D_t^P).$$

*More generally, under the hypotheses of Theorem 3.2 and Corollary 3.5, sufficiently extended inference data can exclude any single static FNN, and indeed any finite family of static FNNs, from the admissible model class.*

*Proof.* For the first claim, let $S$ have finite window size $m$. By Theorem 6.1, any localized perturbation at time $k$ must be forgotten by $S$ after at most $m$ further steps, so the trace predicted by $S$ cannot continue to reflect that perturbation at times $t \geq k+m+1$. By hypothesis, however, the observed trace does exhibit persistence of the perturbation through some such time $t$. Hence the trace predicted by $S$ does not agree with the observed trace through horizon $t$, and therefore

$$S \notin \mathcal{C}_t^P(D_t^P).$$

The second claim follows from Theorem 3.2, which excludes any single static FNN from tracking all future input–output maps under the stated hypotheses, and from Corollary 3.5, which extends this impossibility to finite families of static FNNs. ■

*This proposition gives the relevant non-triviality condition for the nested consistency sets above: extending the behavioral horizon need not merely preserve consistency, but can strictly reduce the admissible class of internal organizations.*

*At the same time, these results do* not *establish strict dependence in the sense of unique identification. Even after long finite traces have ruled out the relevant static surrogates, multiple recurrent or plastic implementations may remain behaviorally indistinguishable over the observed horizon. This non-uniqueness can arise from hidden-unit relabeling, state-space coordinate changes, differences in internal parameterization, or other mechanistic degeneracies invisible at the level of input–output behavior. Thus the consistency set need not collapse to a singleton; at minimum, obvious observational symmetries need not be resolved by finite input–output data alone.*

**Remark 8.3** (Lenient dependency instantiated)
*The preceding results instantiate the qualitative structure of lenient dependency. They rule out* practical independence*, because richer and more temporally extended behavioral traces can exclude broad classes of candidate internal organizations, including bounded-memory static unfoldings and finite families of fixed feedforward surrogates. But they do not establish* strict dependence*, because finite inference data does not, in general, uniquely determine a single recurrent or plastic implementation. Inference data therefore constrains prediction data without uniquely fixing it, which is precisely the intermediate evidential structure emphasized by Kleiner and Hoel.*

[6] *Chang-han Song, Geon Hwang, Jun-hwa Lee, and Myungjoo Kang. Minimal width for universal property of deep rnn.* Journal of Machine Learning Research, *24, 2023.*

[7] *Johannes Kleiner and Erik Hoel. Falsification and consciousness.* Neuroscience of Consciousness, *2021(1):niab001, 04 2021. doi: 10.1093/nc/niab001. URL* https://doi.org/10.1093/nc/niab001.
